# A simple proposal for the publication of journal citation distributions

**DOI:** 10.1101/062109

**Authors:** Vincent Larivière, Véronique Kiermer, Catriona J. MacCallum, Marcia McNutt, Mark Patterson, Bernd Pulverer, Sowmya Swaminathan, Stuart Taylor, Stephen Curry

**Affiliations:** Associate Professor of Information Science, École de bibliothéconomie et des sciences de l’information, Université de Montréal, C.P. 6128, Succ. Centre-Ville, Montréal, QC. H3C 3J7, Canada; Observatoire des Sciences et des Technologies (OST), Centre Interuniversitaire de Recherche sur la Science et la Technologie (CIRST), Université du Québec à Montréal, CP 8888, Succ. Centre-Ville, Montréal, QC. H3C 3P8, Canada; Executive Editor, PLOS, 1160 Battery Street, San Francisco, CA 94111, USA; Advocacy Director, PLOS, Carlyle House, Carlyle Road, Cambridge CB4 3DN, UK; Editor-in-Chief, Science journals, American Association for the Advancement of Science, 1200 New York Avenue, NW, Washington, DC 20005, USA; Executive Director, eLife, 24 Hills Road, Cambridge CB2 1JP, UK; Chief Editor, The EMBO Journal, Meyerhofstrasse 1,69117 Heidelberg, Germany; Head of Editorial Policy, Nature Research, Springer Nature, 225 Bush Street, Suite 1850, San Francisco 94104, USA; Publishing Director, The Royal Society, 6-9 Carlton House Terrace, London SW1Y 5AG, UK; Professor of Structural Biology, Department of Life Sciences, Imperial College, Exhibition Road, London, SW7 2AZ, UK

**Author notes:** Present address: National Academy of Sciences, 2101 Constitution Avenue NW, Washington, DC 20418, USA. Competing interests: The authors have declared that no competing interests exist. Funding: The authors have no funding or support to report.

## Abstract

Although the Journal Impact Factor (JIF) is widely acknowledged to be a poor indicator of the quality of individual papers, it is used routinely to evaluate research and researchers. Here, we present a simple method for generating the citation distributions that underlie JIFs. Application of this straightforward protocol reveals the full extent of the skew of these distributions and the variation in citations received by published papers that is characteristic of all scientific journals. Although there are differences among journals across the spectrum of JIFs, the citation distributions overlap extensively, demonstrating that the citation performance of individual papers cannot be inferred from the JIF. We propose that this methodology be adopted by all journals as a move to greater transparency, one that should help to refocus attention on individual pieces of work and counter the inappropriate usage of JIFs during the process of research assessment.

## Introduction

The problem of over-reliance on the Journal Impact Factor (JIF)^1^ for research and researcher assessment has grown markedly in the 40 years since its original conception in 1972 as a tool for librarians in making decisions on the purchase of journal subscriptions (1). Many stakeholders in academia and academic publishing have recognized that JIFs exert an undue influence in judgements made about individual researchers and individual research papers (2–5).

The main deficiencies of the JIF have been discussed in detail elsewhere (2, 3, 6, 7) but may be summarized as follows: the JIF is calculated inappropriately as the arithmetic mean of a highly skewed distribution of citations^2^; it contains no measure of the spread of the distribution; it obscures the high degree of overlap between the citation distributions of most journals; it is not reproducible and the data that support it are not publicly available (8, 9); it is quoted to a higher level of precision (three decimal places) than is warranted by the underlying data; it is based on a narrow two-year time window that is inappropriate for many disciplines and takes no account of the large variation in citation levels across disciplines (10); it includes citations to ‘non-citable’ items, and citations to primary research papers are conflated with citations to reviews – making the JIF open to gaming and subject to negotiation with Thomson Reuters (7, 11, 12); its relationship with citations received by individual papers is questionable and weakening (13).

We welcome the efforts of others to highlight the perturbing effects of JIFs on research assessment (notably, the San Francisco Declaration on Research Assessment (DORA) (14), the Leiden Manifesto (15), and the Metric Tide report (16)) – and to call for concrete steps to mitigate their influence. We also applaud public statements by funders around the world (*e.g*. Research Councils UK (17), the Wellcome Trust (18), the European Molecular Biology Organisation (EMBO) (19), the Australian Research Council (20), and the Canadian Institutes of Health Research (21)) that no account should be taken of JIFs in assessing grant applications. And we are encouraged by those journals that have cautioned against the misappropriation of JIFs in researcher assessment (7, 11, 22–25).

At the same time we recognize that many academics and many institutions lack confidence in the ability of the members of funding, promotion or other research assessment panels to shed what has become a habit of mind. This is exacerbated by the fact that various quantitative indicators are increasingly part of the toolbox of research management (16) and are often viewed as a convenient proxy for ‘quality’ by busy academics perennially faced with sifting through large numbers of grant applications or CVs.

To challenge the over-simplistic interpretation of JIFs, we present here a straightforward methodology for generating the citation distribution of papers published in any journal. Consistent with previous analyses (9, 26), application of this method to a selection of journals covering a number of different scientific disciplines shows that their citation distributions are skewed such that most papers have fewer citations than indicated by the JIF and that the spread of citations per paper typically spans two to three orders of magnitude resulting in a great deal of overlap in the distributions for different journals. Although these features of citation distributions are well known to bibliometricians and journal editors (7, 23, 26), they are not widely appreciated in the research community. It is the desire to broaden this awareness that motivated us, a group drawn from the research, bibliometrics and journals communities, to conduct the analysis reported here.

We believe that the wider publication of citation distributions provides a healthy check on the misuse of JIFs by focusing attention on their spread and variation, rather than on single numbers that conceal these universal features and assume for themselves unwarranted precision and significance. We propose that this methodology be adopted by all journals that publish their impact factors so that authors and readers are provided with a clearer picture of the underlying data. This proposal echoes the reasonable requests that journal reviewers and editors make of authors to show their data in justifying the claims made in their papers.

## Methods

### Purchased Database Method

The analyses presented here were conducted using the three main citation indexes purchased from Thomson Reuters by the Observatoire des sciences et des technologies (OST-UQAM): the Science Citation Index Expanded, the Social Science Citation Index, and the Arts and Humanities Citation index. Data were obtained on March 18^th^ 2016 and the results reflect the content of the database at that point in time. They may therefore differ from results obtained subsequently using its Web version, the Web of Science^™^, which is continuously updated (see below), though any differences are likely to be small for distributions calculated over equivalent time windows.

To obtain the number of citations per citable item (which we defined as articles and reviews, following Thomson Reuters practice in JIF calculations (27)), we used Thomson Reuters’ matching key to define links between citing and cited papers. As part of our analysis, additional citations were retrieved from the database using the various forms of each journal’s name^3^. Although these could not be linked to specific papers and cannot therefore be included in the citation distributions, they are listed as unmatched citations in Table 1 to give an idea of the numbers involved. It is worth noting that these unmatched citations are included in the calculation of the JIF. For the journals *eLife*, *Scientific Reports, Proceedings of the Royal Society B: Biological Sciences*, and *Nature Communications*, the share of unmatched citations is higher, which suggests that citations to specific papers are underestimated by the Thomson Reuters matching key (Table 1). Thus, these distributions underestimate the numbers of citations per paper — and may overestimate the numbers of papers with zero citations. Given that these unmatched citations are likely to be evenly distributed across all papers, this effect should not affect the structure of the distributions.

**Table 1.** Citations received in 2015 by document type published in 2013 and 2014.

| Journal | Article |  | Review |  | Correction |  | Editorial-Material |  | Others documents |  | Unmatched Citations |  | Total Citations |
| --- | --- | --- | --- | --- | --- | --- | --- | --- | --- | --- | --- | --- | --- |
|  | N. | % | N. | % | N. | % | N. | % | N. | % | N. | % |  |
| eLife | 5,459 | 84.4% |  |  | 10 | 0.2% | 98 | 1.5% |  |  | 902 | 13.9% | 6,469 |
| EMBO J. | 3,219 | 82.2% | 472 | 12.1% | 2 | 0.1% | 121 | 3.1% | 4 | 0.1% | 97 | 2.5% | 3,915 |
| J. Informetrics | 387 | 92.6% | 6 | 1.4% | 1 | 0.2% |  |  | 10 | 2.4% | 14 | 3.3% | 418 |
| Nature | 54,143 | 83.2% | 3,554 | 5.5% | 47 | 0.1% | 2,770 | 4.3% | 1,681 | 2.6% | 2,903 | 4.5% | 65,098 |
| Nature Comm. | 43,957 | 88.5% | 82 | 0.2% | 15 | 0.0% |  |  |  |  | 5,609 | 11.3% | 49,663 |
| PLOS Biol. | 2,927 | 87.0% | 16 | 0.5% |  |  | 201 | 6.0% |  |  | 219 | 6.5% | 3,363 |
| PLOS Genet. | 9,964 | 91.6% | 238 | 2.2% | 3 | 0.0% | 46 | 0.4% |  |  | 621 | 5.7% | 10,872 |
| PLOS ONE | 168,590 | 90.7% | 2,753 | 1.5% | 86 | 0.0% | 5 | 0.0% |  |  | 14,378 | 7.7% | 185,812 |
| Proc. R. Soc. B | 4,462 | 76.3% | 436 | 7.5% | 4 | 0.1% | 31 | 0.5% |  |  | 916 | 15.7% | 5,849 |
| Science | 43,665 | 75.6% | 5,816 | 10.1% | 4 | 0.0% | 4,522 | 7.8% | 1,011 | 1.8% | 2,747 | 4.8% | 57,765 |
| Sci. Rep. | 29,668 | 86.2% | 1 | 0.0% | 11 | 0.0% | 2 | 0.0% |  |  | 4,750 | 13.8% | 34,432 |

### Subscription Database Method

The use of a purchased database provides convenient access the bulk citation data, but the expense involved means the method described above is only likely to be a viable option for professional bibliometricians. To facilitate the generation of citation distributions by non-specialists, we developed step-by-step protocols that rely on access to essentially the same data via subscription to either the Web of Science^™^ (Thomson Reuters Inc.) or Scopus^™^ (Elsevier BV). The details of each protocol are presented in Appendices 1 and 2^4^.

It should be noted that all the protocols we present here for generating distributions use only those citations that are unambiguously matched to specific papers. This is in contrast to the approach used by Thomson Reuters in calculating JIFs which includes citations to all document types as well as unmatched citations (see Table 1). Thus, while the cohort of articles can be matched to the JIF cohort (namely, citations received in 2015 to articles published in 2013 and 2014) the absolute values of the citations to individual articles and the total number of citations can vary substantially from that used in the JIF calculation.

### Data presentation

The appearance of citation distributions will vary depending on the choice of vertical scales and the degree of binning used to aggregate citation counts. We have opted to use vertical scales that vary to reflect the different publishing volumes of the journals included in our study. We have also chosen not to bin the citation counts except for papers that have 100 or more citations. These choices maximize the resolution of the plots while restricting them to a common horizontal scale that facilitates comparison of the shapes of the distributions (see Fig. 1) – and we would recommend them as standard practice. For more direct comparisons between journals, the distributions can be replotted with a common vertical scale (*e.g*. using percentages, as in Fig. 4b).

**Fig 1.**
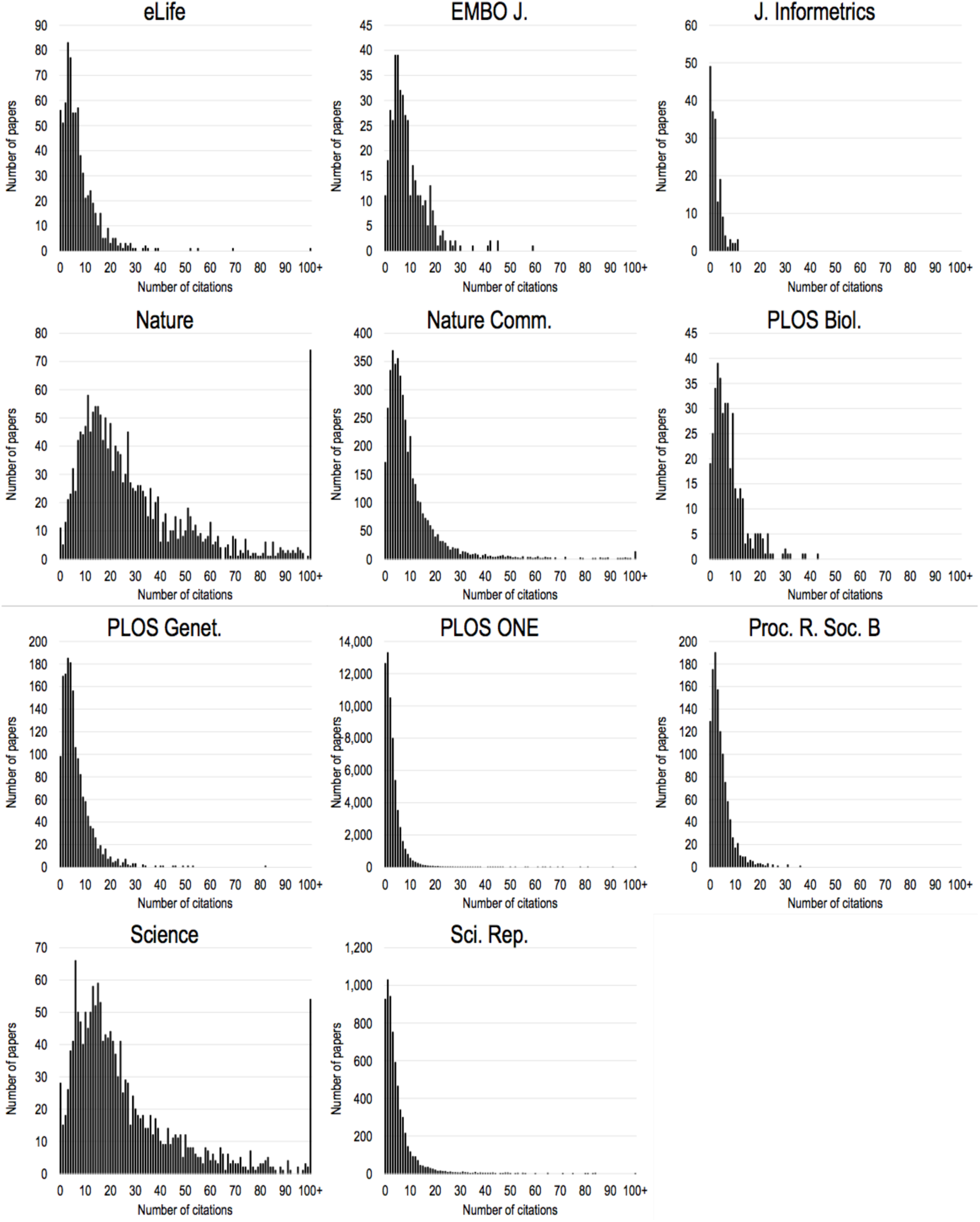
Citation distributions of 11 different science journals. Citations are to ‘citable documents’ as classified by Thomson Reuters, which include standard research articles and reviews. The distributions contain citations accumulated in 2015 to citable documents published in 2013 and 2014 in order to be comparable to the 2015 JIFs published by Thomson Reuters. To facilitate direct comparison, distributions are plotted with the same range of citations (0-100) in each plot; articles with more than 100 citations are shown as a single bar at the right of each plot.

## Results

Using the Purchased Database Method described above, we generated frequency plots – or citation distributions – for 11 journals: *eLife*, *EMBO Journal, Journal of Informetrics*, *Nature*, *Nature Communications*, *PLOS Biology*, *PLOS Genetics*, *PLOS ONE*, *Proceedings of the Royal Society B: Biological Sciences*, *Science*, and *Scientific Reports* (Figure 1). The journals selected are both multidisciplinary and subject-specific in scope, and range in impact factor from less than 3 to more than 30. They represent journals from seven publishers: the American Association for the Advancement of Science (AAAS), eLife Sciences, Elsevier, EMBO Press, Springer Nature, the Public Library of Science (PLOS), and the Royal Society.

In an attempt to relate our analyses to the widely-available JIFs for 2015, the period over which the citations accumulated for our distributions was chosen to match that of the 2015 Journal Impact Factors published by Thomson Reuters – namely, the number of citations accrued in 2015 from documents published in 2013-2014. However, to more effectively compare journal distributions, we opted to include only citable items as classified by Thomson Reuters, which includes standard research articles and review articles (27), because different journals publish different amounts of additional content such as editorials, news items, correspondence, and commentary. It should also be noted that the definition of research and review articles used by Thomson Reuters does not always match the labels given to different document types by journals. Table 1 provides a summary of the number and percentage of articles and citations accrued for each document type within each journal as classified by Thomson Reuters. The summary data used to generate the distributions are provided in Supplemental File 1.

While the distributions presented in Figure 1 were generated using purchased data (see Methods), we tested whether similar distributions could be produced following the step-by-step Subscription Based Method outlined in Appendix 1 which uses data accessed online via Web of Science™. As seen in the distributions calculated for the EMBO Journal (Figure 2), the broad features of the distributions from these different sources are essentially identical, with differences being due to updates made on the database between purchase of data and time of online access.

**Fig 2.**
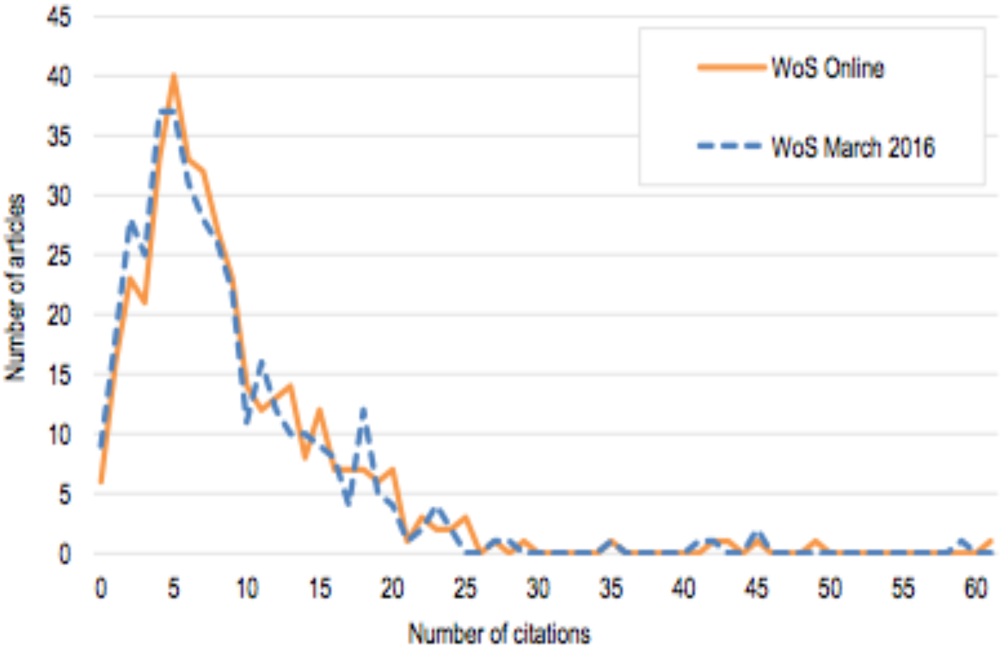
Comparison plot for EMBO Journal. The analyses in this paper are based on proprietary data bought from Thomson Reuters by the Observatoire des sciences et des technologies (OST-UQAM) and is similar to that used by Thomson Reuters to generate the JIFs (‘WoS March 2016’). Publishers and Institutions with a subscription to the Web of Science™ have access to a different dataset (‘WoS online’).

For all journals, the shape of the distribution is highly skewed to the right, the left-hand portion being dominated by papers with lower numbers of citations. Typically, 65-75% of the articles have fewer citations than indicated by the JIF (Table 2). The distributions are also characterized by long rightward tails; for the set of journals analyzed here, only 15-25% of the articles account for 50% of the citations as shown in the cumulative distributions plotted in Figure 3. The distributions are also broad, often spanning two or more orders of magnitude. The spread tends to be broader for journals with higher impact factors. Our results also show that journals with very high impact factors tend to have fewer articles with low numbers of citations.

**Table 2:** Percentage of papers published in 2013-2014 with number of citations below the value of the 2015 JIF.

| Journal | JIF | % citable items below JIF |
| --- | --- | --- |
| eLife | 8.3 | 71.2% |
| EMBO J. | 9.6 | 66.9% |
| J. Informetrics | 2.4 | 68.4% |
| Nature | 38.1 | 74.8% |
| Nature Comm. | 11.3 | 74.1% |
| PLOS Biol. | 8.7 | 66.8% |
| PLOS Genet. | 6.7 | 65.3% |
| PLOS ONE | 3.1 | 72.2% |
| Proc. R. Soc. B | 4.8 | 65.7% |
| Science | 34.7 | 75.5% |
| Sci. Rep. | 5.2 | 73.2% |

**Fig 3.**
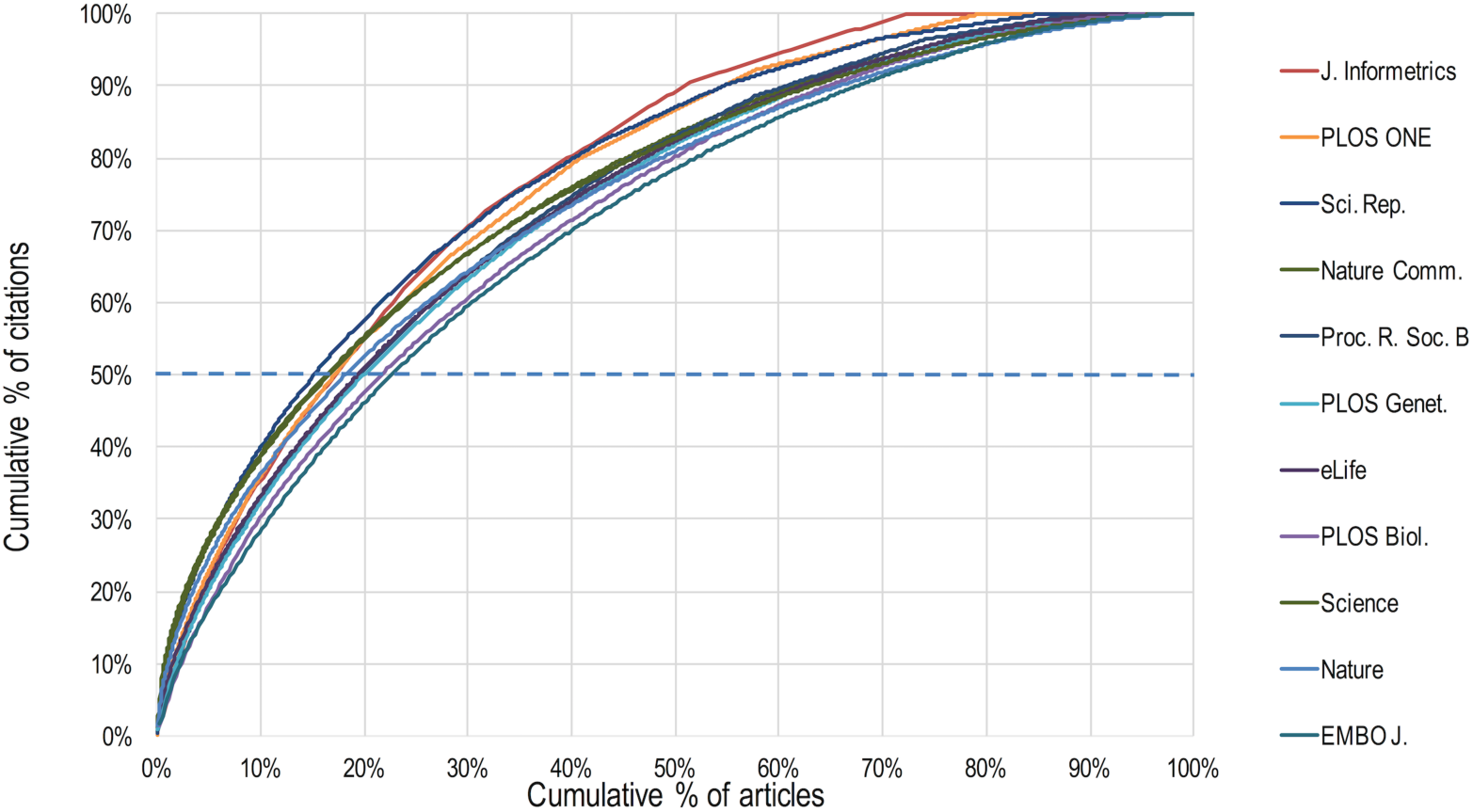
The cumulative % of citations and articles plotted for the 11 journals included in this study. The plots for all the journals are very similar, which reflects the skewness of the distributions shown in Figure 1.

The journals with highest impact factors (*Nature* and *Science*) also tend to have more articles with very high levels of citation within the two-year time period used for JIF calculations (and our analyses). The most cited articles in *Nature* and *Science* are cited 905 times and 694 times respectively in 2015 (see Supplemental File 1). Highly cited articles also appear in journals with much lower impact factors; for example, the most-cited articles in *PLOS ONE* and *Scientific Reports* are cited 114 and 141 times in 2015, respectively. For all journals, the very highly cited articles represent a small percentage of the total number of articles and yet have a disproportionate influence on the impact factor because it is based on an arithmetic mean calculation that does not take proper account of the skew in the distribution.

Despite the variations in citation distributions between journals that are evident in Figure 1, there is substantial overlap in the citation distributions across all the journals (Figure 4a). The overlap becomes more apparent when the number of articles are converted to a percentage which has the effect of placing the data on a common vertical scale (Figure 4b). This makes it clear that, even without taking into account the effect of the sizes of different disciplines on citation counts, papers with high and low numbers of citations appear in most, if not all, journals.

**Fig 4.**
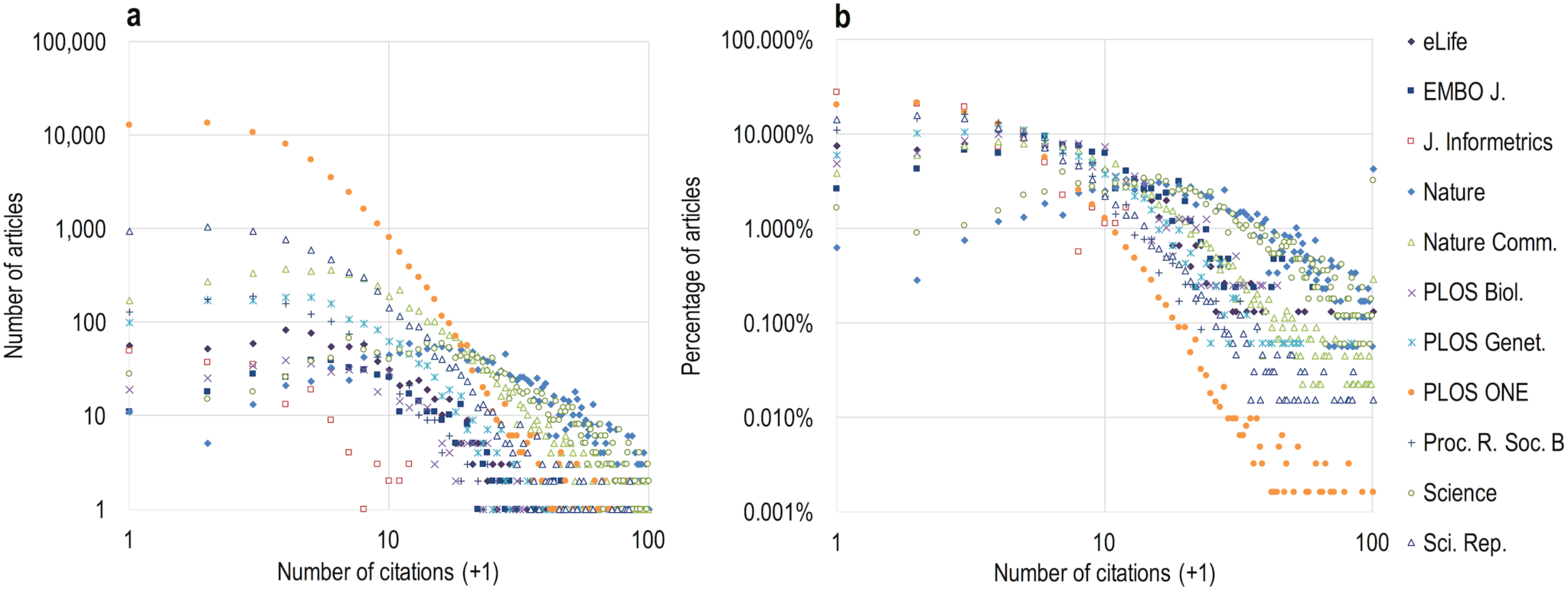
A log-scale comparison of the 11 citation distributions. (a) The absolute number of articles plotted against the number of citations. (b) The percentage of articles plotted against the number of citations.

## Discussion

The aim of this paper is to increase awareness of the journal citation distributions underlying JIFs by disseminating a simple protocol that allows them to be produced by anyone with access, via institutional or publisher subscription, to Web of Science™ or Scopus™ (Appendices 1 and 2). We have selected a group of journals for illustrative purposes and have made no attempt to be comprehensive. Our intention here is to encourage publishers, journal editors and academics to generate and publish journal citation distributions as a countermeasure to the tendency to rely unduly and inappropriately on JIFs in the assessment of research and researchers.

The proposed method is straightforward and robust. It yields citation distributions that have all the same features identified in previous analyses (9, 26). The distributions reveal that, for all journals, a substantial majority of papers have many fewer citations than indicated by the arithmetic mean calculation used to generate the JIF, and that for many journals the spread of citations per paper varies by more than two orders of magnitude. Although JIFs do vary from journal to journal, the most important observation as far as research assessment is concerned, and one brought to the fore by this type of analysis, is that there is extensive overlap in the distributions for different journals. Thus for all journals there are large numbers of papers with few citations and relatively few papers with many citations.

Arguably, an alternative to publishing citation distributions might be for journals to provide additional metrics or parameters to describe the distributions (such as the median, skew or inter-quartile range), to give readers a clearer idea of the variation in the citations to the papers therein (7, 28). While such additional information may indeed be useful, we think that the transparency provided by presenting the full distribution and showing the noise inherent in the data give a more powerful visual representation of the underlying complexity.

In our view, the variation evident in the distributions underscores the need to examine each paper on its own merits and serves as a caution against over-simplistic interpretations of the JIF. We do not wish to underestimate the difficulties inherent in making a fair assessment of a paper or a researcher’s publication record. The issue is complicated because journal prestige is often elided with the JIF, and is one of several factors (*e.g*. reputation, audience, format) that influence authors’ choices of publishing venues for their own work. The name of the journal where a paper is published and the numbers of citations that it (or the journal) attracts might reasonably be thought of as interesting pieces of information in assessing a piece of work, but we would argue strongly that the process of assessment has to go beyond mere branding and numbers. Users of JIFs (or indeed users of citation counts – since there are many reasons for citing a paper) should appreciate these and other complicating factors, such as the inflationary effect on citations in journals with higher JIFs, which may be due to greater visibility and perceived prestige (29–31). This effect is illustrated by analysis of citations to a medical “white paper” that was published in eight different journals in 2007 and showed that the number of citations that each publication received correlated strongly (R^2^ = 0.91) with the JIF of the host journal across a range of JIF values from 2 to 53 (32).

With one exception (*J. Informetrics*), our analyses cover a collection of journals that are generally broad in scope, encompassing several different disciplines across the sciences. It may be that the breadth of the distribution is less marked in journals of narrower scope, although their JIFs are just as prone to outlier effects, and overlapping distributions of citations have been observed in more specialized journals (9, 33).

Despite the overlap, there are evident differences in the *average* citation performance of different journals, and we are not arguing that the JIF is of no interest in comparing different journals (though such comparisons should take account Royle’s analysis of significance of differences between JIFs (9)). However, our primary interest here is not to compare journals but to mitigate the problem of applying journal-based metrics in *individualized* research assessment. The properties of the distributions help to highlight how difficult it is to extract reliable citation information about a given paper from the JIF – as demonstrated recently by Berg’s analysis of our distribution data (34) – never mind the trickier business of assessing its *quality*. The aim of publishing citation distributions is to expose the exaggerated value often attributed to the JIF and thereby strengthen the contention that it is an indicator that is all too easily misconstrued in the evaluation of research or researchers.

On a technical point, the many unmatched citations (*i.e*. citations not clearly linked to a specific article, Table 1) that were discovered in the data for *eLife*, *Nature Communications*, *Proceedings of the Royal Society: Biological Sciences*, and *Scientific Reports* raises concerns about the general quality of the data provided by Thomson Reuters. Searches for citations to *eLife* papers, for example, have revealed that the data in the Web of Science™ are incomplete owing to technical problems that Thomson Reuters is currently working to resolve (35). We have not investigated whether similar problems affect journals outside the set used in our study and further work is warranted. However, the raw citation data used here are not publicly available but remain the property of Thomson Reuters. A logical step to facilitate scrutiny by independent researchers would therefore be for publishers to make the reference lists of their articles publicly available. Most publishers already provide these lists as part of the metadata they submit to the Crossref metadata database (36). They can easily permit Crossref to make them public, though relatively few have opted to do so. If all Publisher and Society members of Crossref (over 5,300 organisations) were to grant this permission, it would enable more open research into citations in particular and into scholarly communication in general (36).

## Conclusion and Recommendations

The co-option of JIFs as a tool for assessing individual articles and their authors, a task for which they were never intended, is a deeply embedded problem within academia. There are no easy solutions. It will not suffice to change the way that citation statistics are presented. We recognize alongside De Rijcke and Rushforth (37) that the roots of the problem need to be addressed upstream, and not simply post-publication. Nevertheless, we hope that by facilitating the generation and publication of journal citation distributions, the influence of the JIF in research assessment might be attenuated, and attention focused more readily onto the merits of individual papers, and onto the diverse other contributions of researchers such as sharing data, code, and reagents (not to mention their broader – such as peer review and mentoring students – to the mission of the academy). We would also encourage ongoing efforts to diversify assessment procedures in practical ways (*e.g*. through the use of summaries of the researcher’s key publications and achievements) and to address the particular challenges of assessing interdisciplinary research (38, 39).

To advance this agenda we therefore make the following specific recommendations:

- *We encourage journal editors and publishers that advertise or display JIFs to publish their own distributions using the above method, ideally alongside statements of support for the view that JIFs have little value in the assessment of individuals or individual pieces of work (see this example at the Royal Society). Large publishers should be able to do this through subscriptions to Web of Science™ or Scopus™; smaller publishers may be able to ask their academic editors to generate the distributions for their journals*.
- *We encourage publishers to make their citation lists open via Crossref, so that citation data can be scrutinized and analyzed openly*.
- *We encourage all researchers to get an ORCID_iD, a digital identifier that provides unambiguous links to published papers and facilitates the consideration of a broader range of outputs in research assessment*.

These recommendations represent small but feasible steps that should improve research assessment. This in turn should enhance the confidence of researchers in judgements made about them and, possibly, the confidence of the public in the judgements of researchers. This message is supported by the adoption in many journals of article-level metrics and other indicators that can help to track the use of research paper within and beyond the academy. We recognize that drawing attention to citation distributions risks inadvertent promotion of JIFs. However, we hope that the broader message is clear: research assessment needs to focus on papers rather than journals, keeping in mind that downloads and citation counts cannot be considered as reliable proxies of the quality of an individual piece of research (16). We would always recommend that a research paper is best judged by reading it.

## Acknowledgements

We are very grateful to Valda Vinson and Monica Bradford for critical reading of the manuscript. We also would like to thank everyone who took the time to comment on the first version of this preprint; most of this commentary can be accessed through bioRxiv. We particularly thank Ludo Waltman and Sarah de Rijcke for critical reading of the revised manuscript.

## Supplemental Files

**Supplemental File 1**: Microsoft Excel spreadsheet containing the summary data used to prepare the Figures and Tables for this paper. Also contains the Figures and Tables themselves.

**Supplemental File 2:** Microsoft PowerPoint file containing ready-to-use, high-resolution slides of the Figures and Tables from this paper.

**Supplemental File 3:** PDF version of Supp. File 2, containing ready-to-use, high-resolution slides of the Figures and Tables from this paper.

**Supplemental File 4:** Responses to Comments on the first version of this preprint (PDF)

## Author Contributions Statement

**Vincent Larivière:** methodology, formal analysis, investigation, writing – original draft preparation, visualization

**Véronique Kiermer:** writing – review and editing

**Catriona MacCallum:** writing – original draft preparation, review and editing

**Marcia McNutt:** writing – review and editing

**Mark Patterson:** writing – original draft preparation, review and editing

**Bernd Pulverer:** writing – review and editing

**Sowmya Swaminathan;** writing – review and editing

**Stuart Taylor:** methodology, formal analysis, investigation, writing

– original draft preparation, visualization

**Stephen Curry:** conceptualization, investigation, writing – original draft preparation, review and editing

## Footnotes

1 *The JIF is formally defined as the mean number of citations received in a given year by papers published in a journal over the two previous years*.

2 *Although the JIF is presented as an arithmetic mean, the numerator is the total number of citations received by all documents published in the journal whereas the denominator is the subset of documents that Thomson Reuters classifies as ‘citable’ (i.e. ‘Articles’ and ‘Reviews’)*.

3 *For example, the journal Proceedings of the Royal Society B – Biological Sciences appeared in the reference list as P R SOC B, P R SOC B IN PRESS, P R SOC BIOL SCI, P R SOC LONDON B, etc*.

4 *Since there are more journals and papers indexed in Scopus^™^, citation rates for individual articles are likely to be higher than those presented here if this database is used to generate distributions*.

## Appendix 1 Method for generating the journal citation distribution graph from the Web of Science^™^ (2014 Impact Factor set)

The example given below is for generating distributions over the two-year window (2012-2013) that is used in calculation of the 2014 Journal Impact Factor. For later years, such as for the distributions based on the 2015 JIF in the main article here, the two-year window should be adjusted accordingly.

1. In <u>Web of Science</u>, select *Core Collection*.

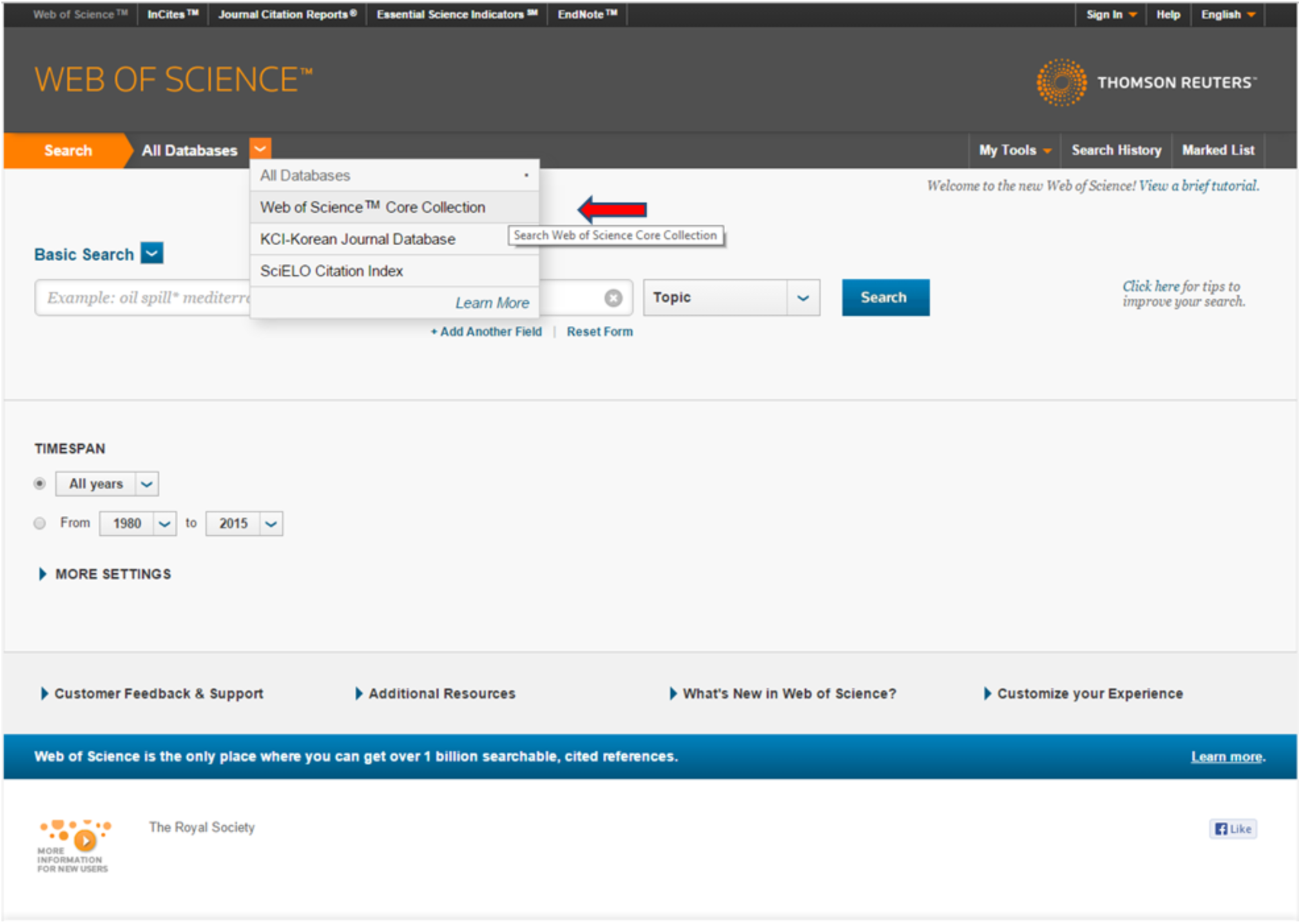

2. Select ‘Publication Name’ as the filter for the first field and then enter the journal name in the associated free text box (alternatively the journal title can be selected from the ‘Select from index’ link). Select the ‘Add Another Field’ option and select ‘Year Published’ as the second filter and enter 2012-2013 in the text box. Click search. In the example shown, the journal *Biology Letters* has been selected.

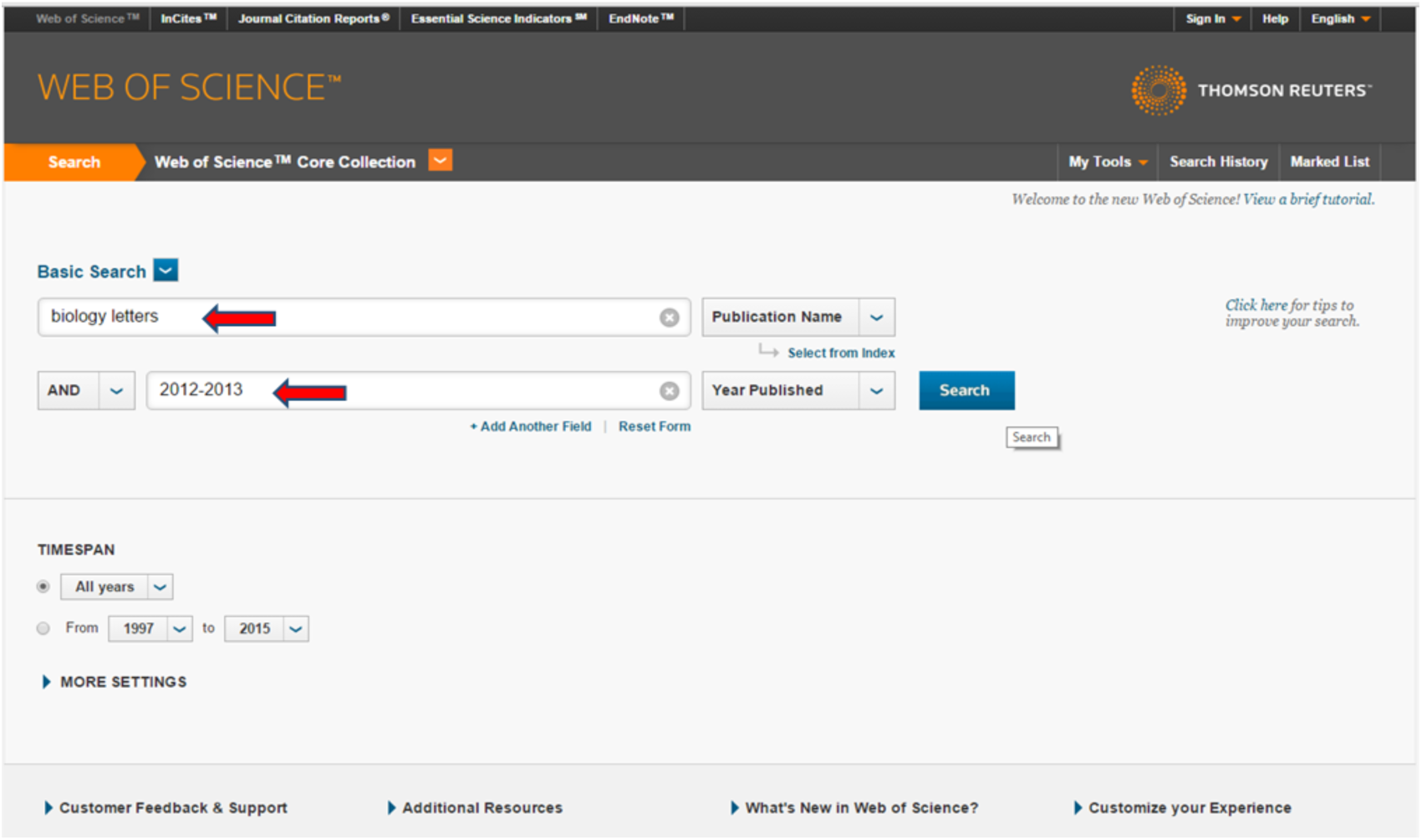

3. That produces the requisite article set. Next, click *Create Citation Report*. (To match as closely as possible the distributions shown in the analyses in this paper, limit the search to ‘Articles’ and ‘Reviews’ using the buttons on the left hand side of the screen under ‘Document Types’.). Note that, as in the screenshot below, if the journal does not publish reviews (as classified by Thomson Reuters), an option to tick ‘Reviews’ will not be available.

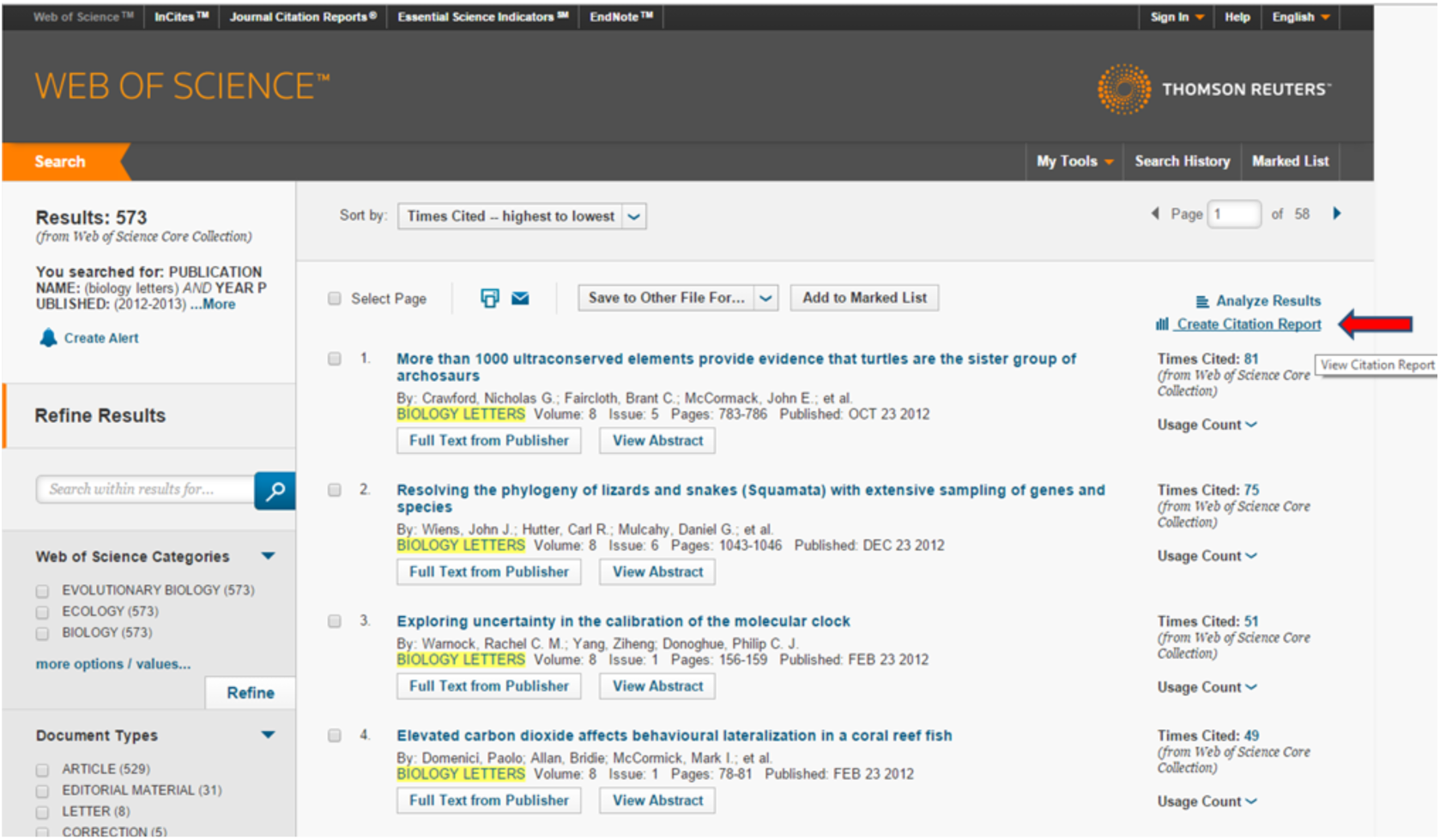

4. The citation report should look similar to this. Note the number of articles retrieved by the search at the top of the page (573 in example below).

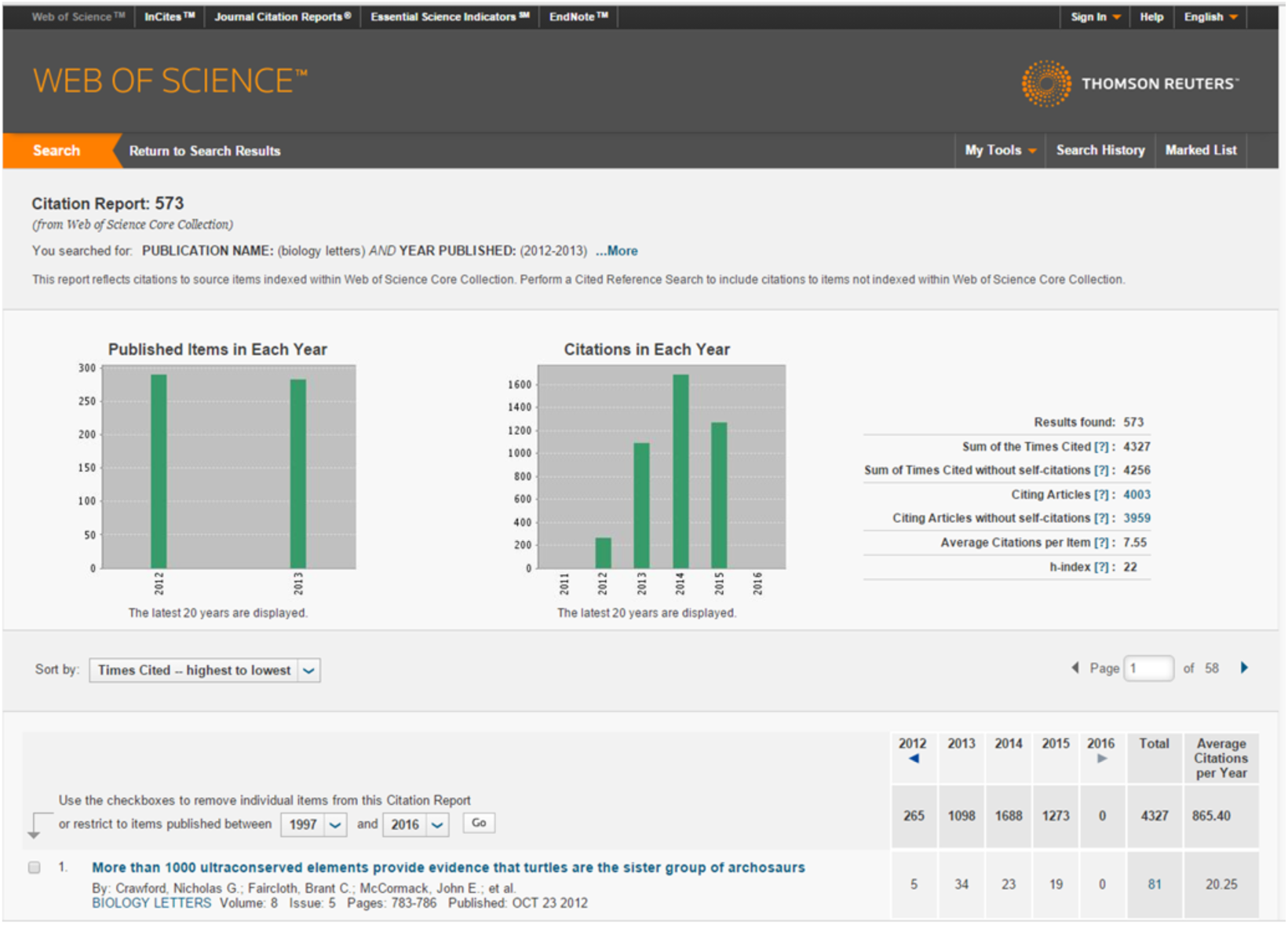

5. Scroll to the bottom of the web-page and export the list to Excel.

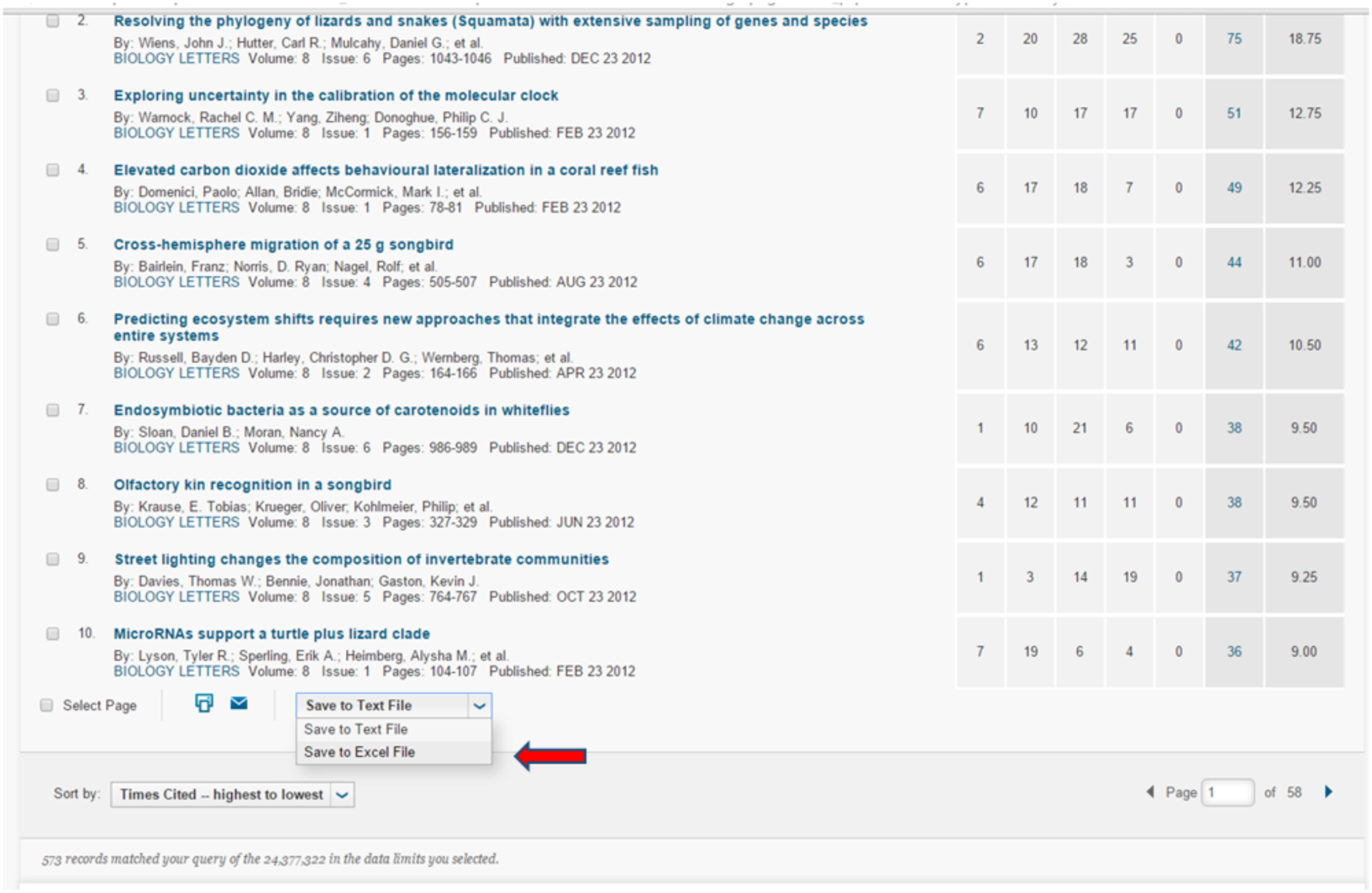

6. When prompted, enter the number of articles retrieved by the search as the maximum number of records. Web of Science^™^ will only process 500 records at a time, so if you have more articles than that, you’ll need to export several Excel files and then combine them.

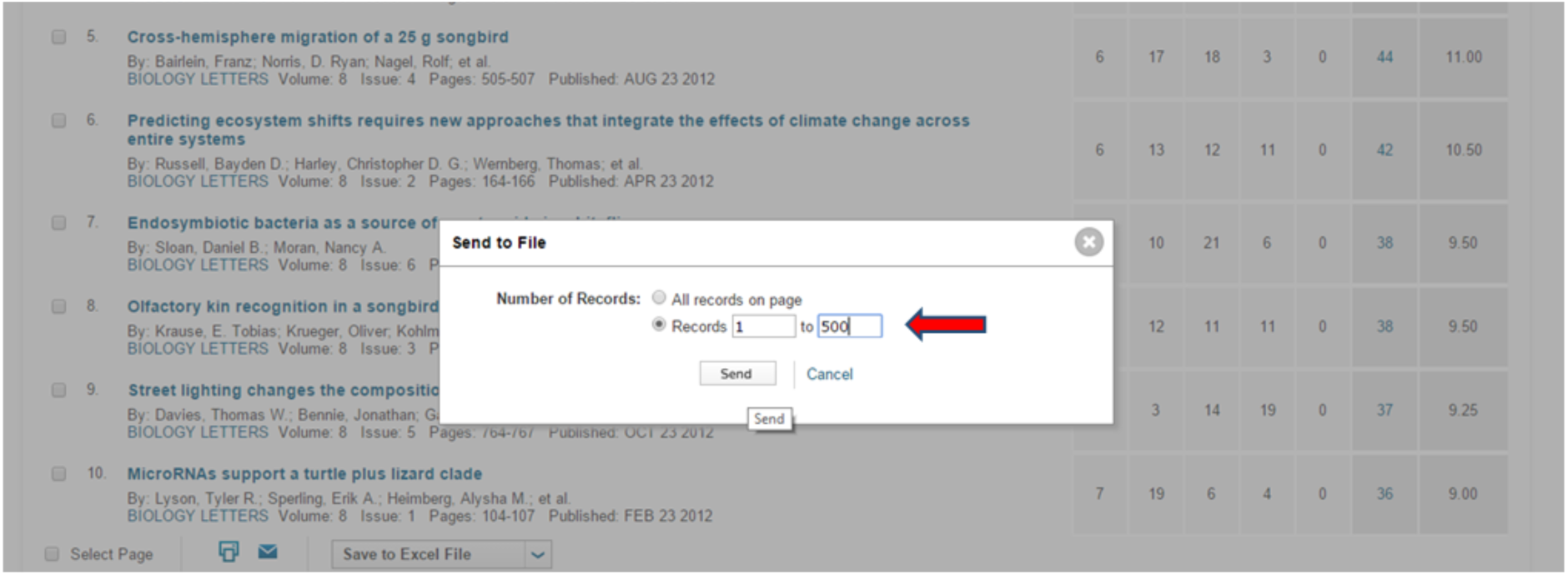

7. Open the combined file in Excel.

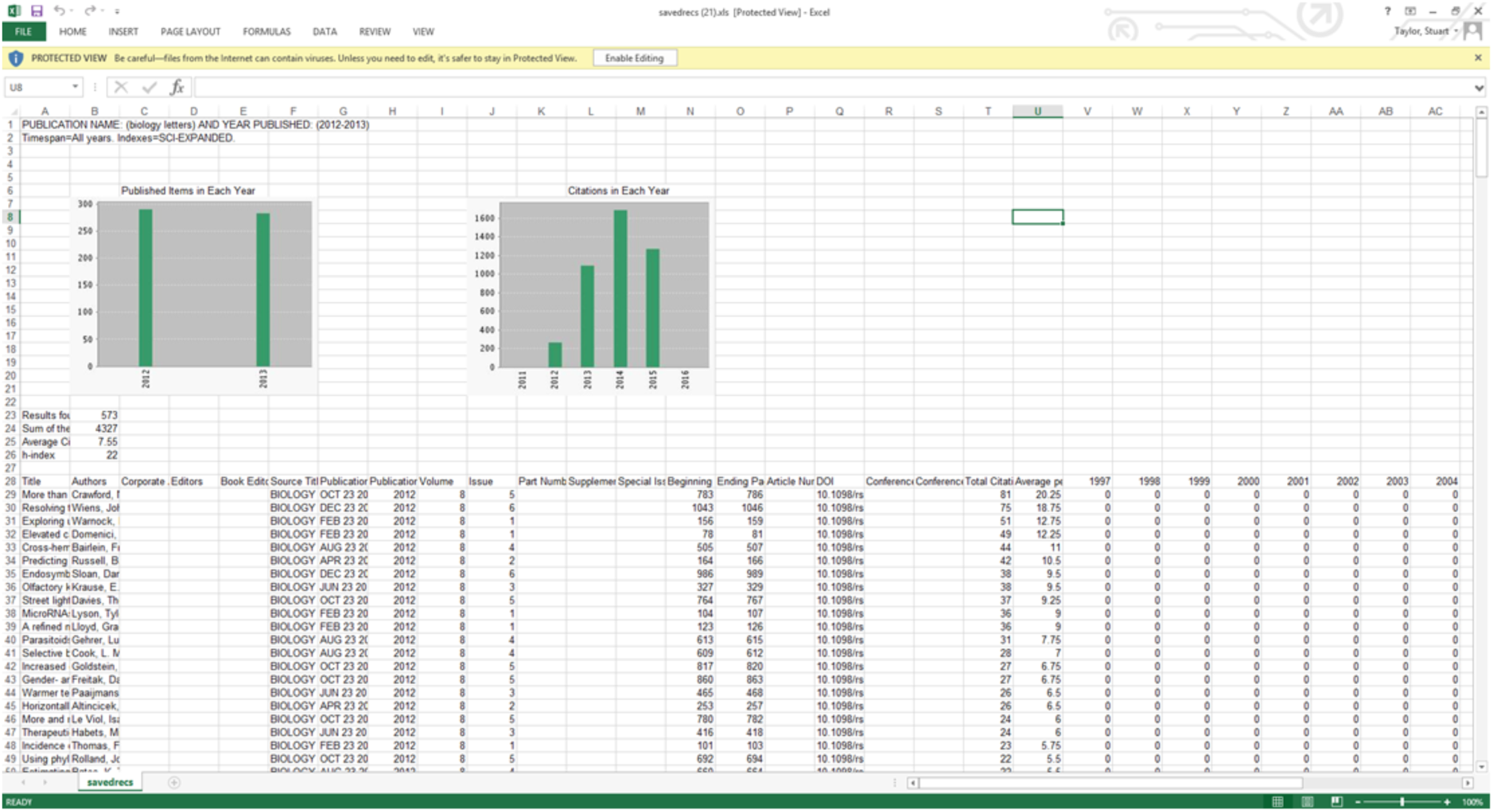

8. Only the column for the citations received in 2014 is needed for the distribution, so scroll across and select that column.

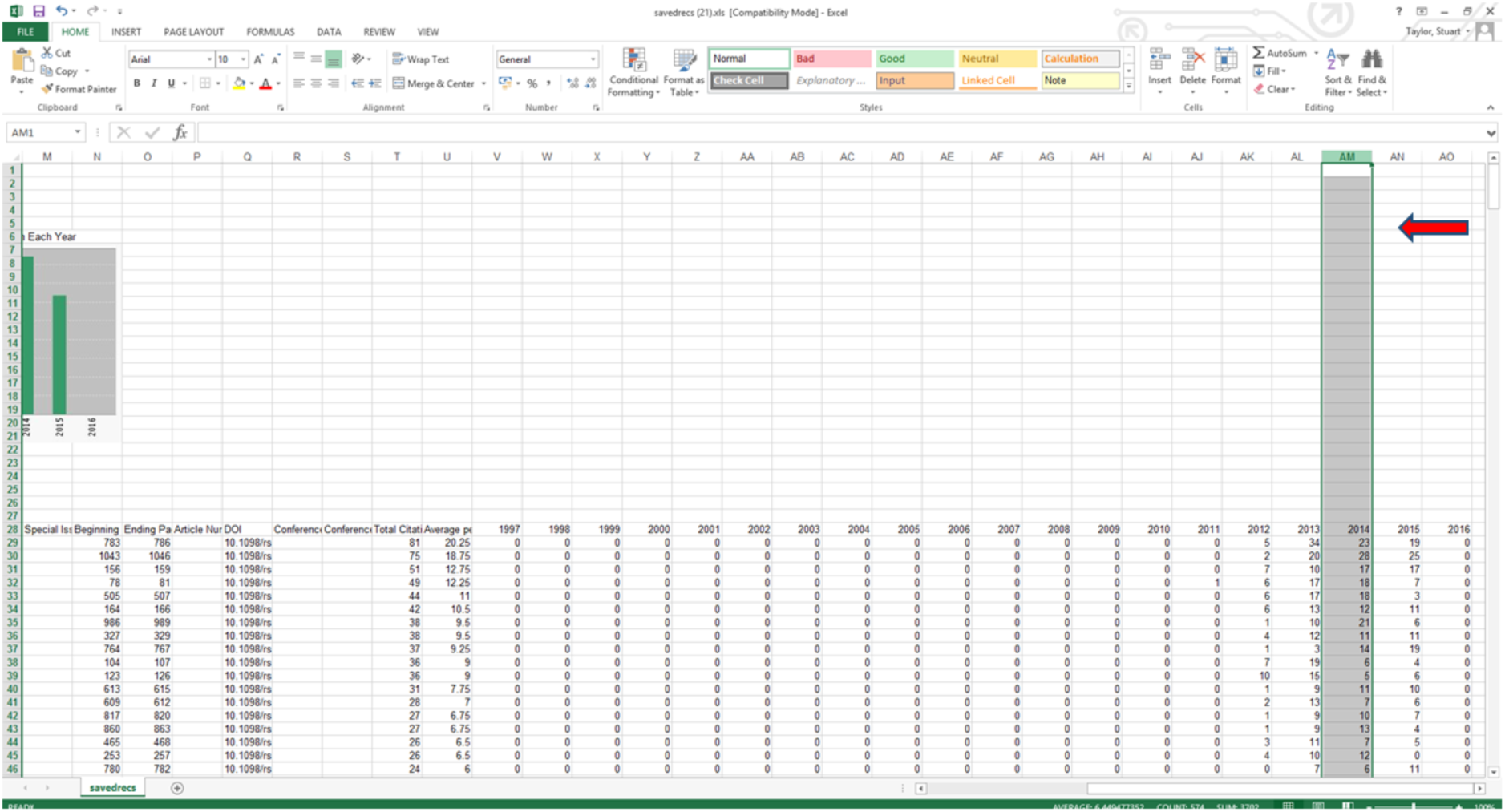

9. Sort the column into descending order (omitting the ‘2014’ label at the top).

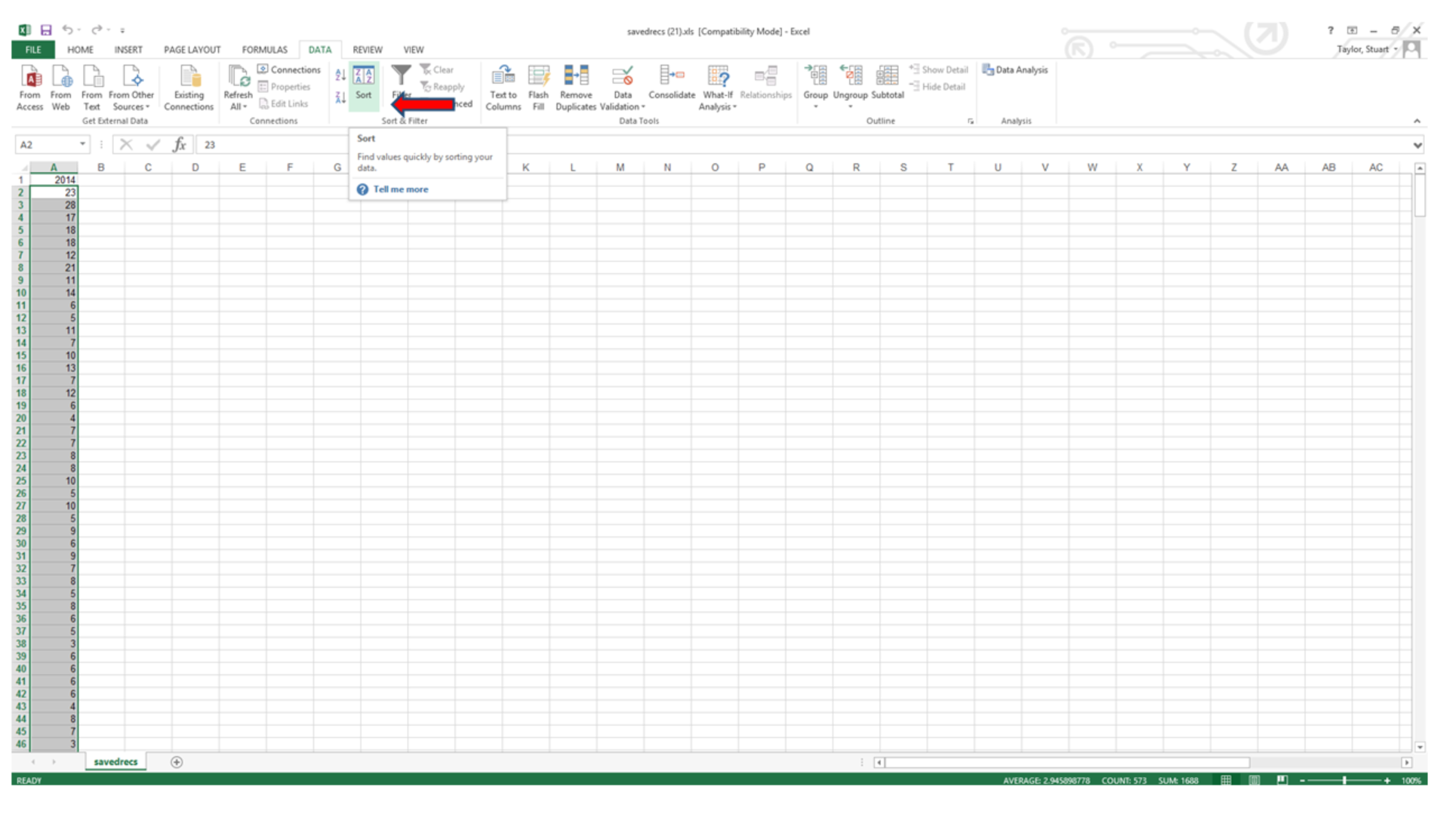

10. Note the maximum citation (x) count and create a new column containing 0 to X called “Citations”. In the example shown below, x = 28.

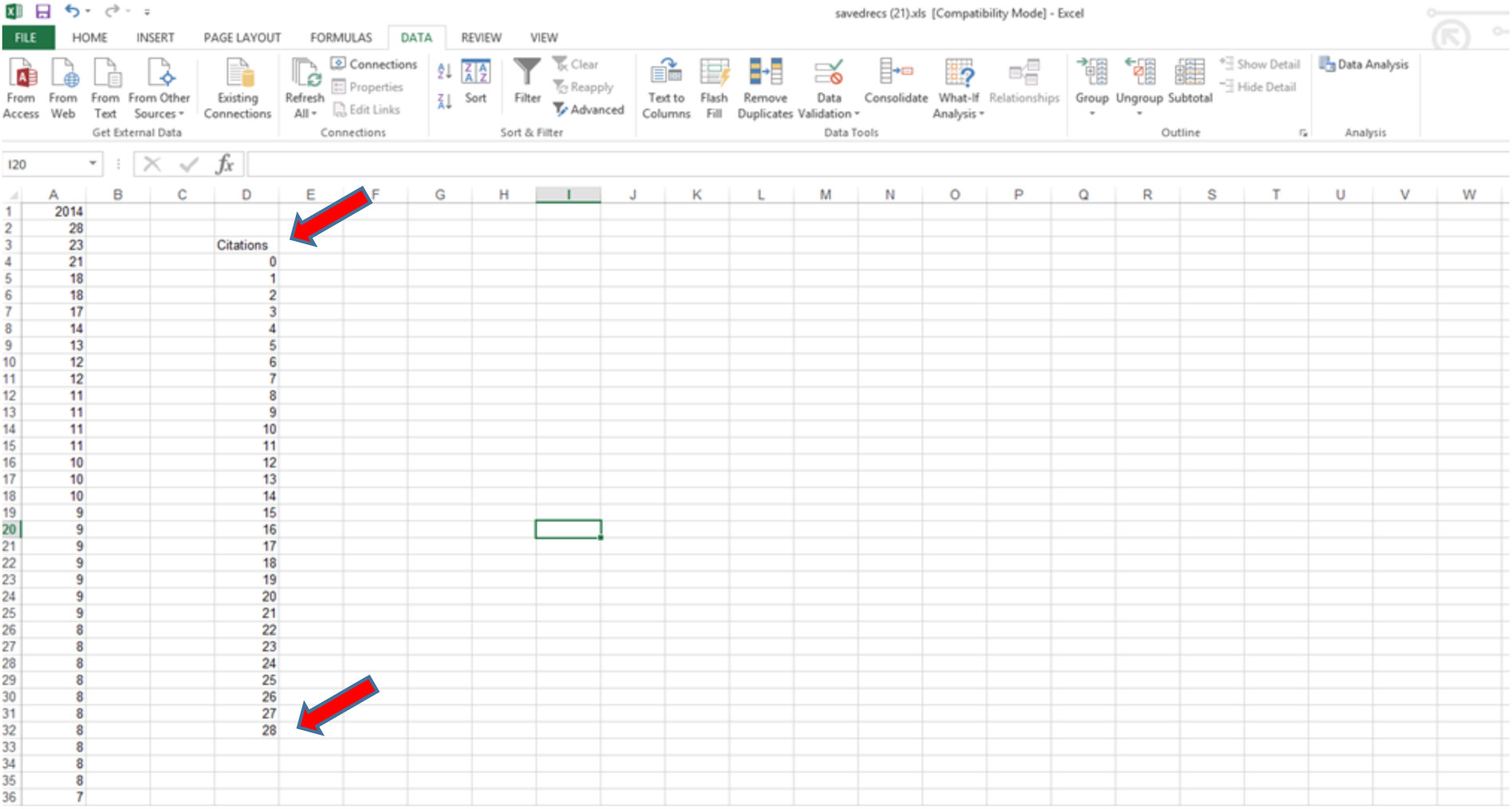

11. Enter the formula =COUNTIF(A:A,D4) into the cell *next* to the 0 citations (where A is the column containing the citations, and D4 is the cell indicating zero citations – see below).

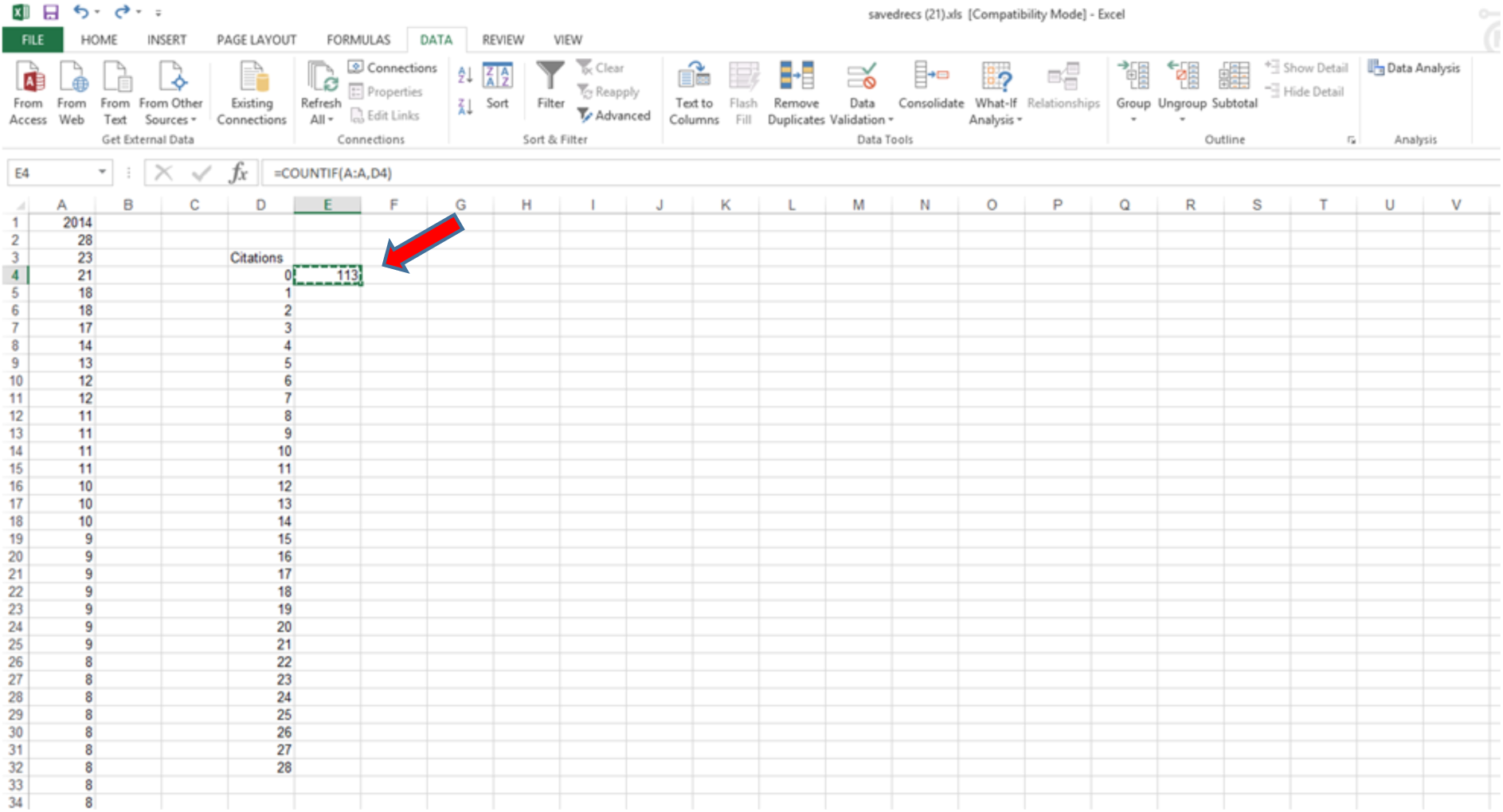

12. Copy and paste this formula into the remaining cells in the Citations column. This generates the data for the frequency distribution.

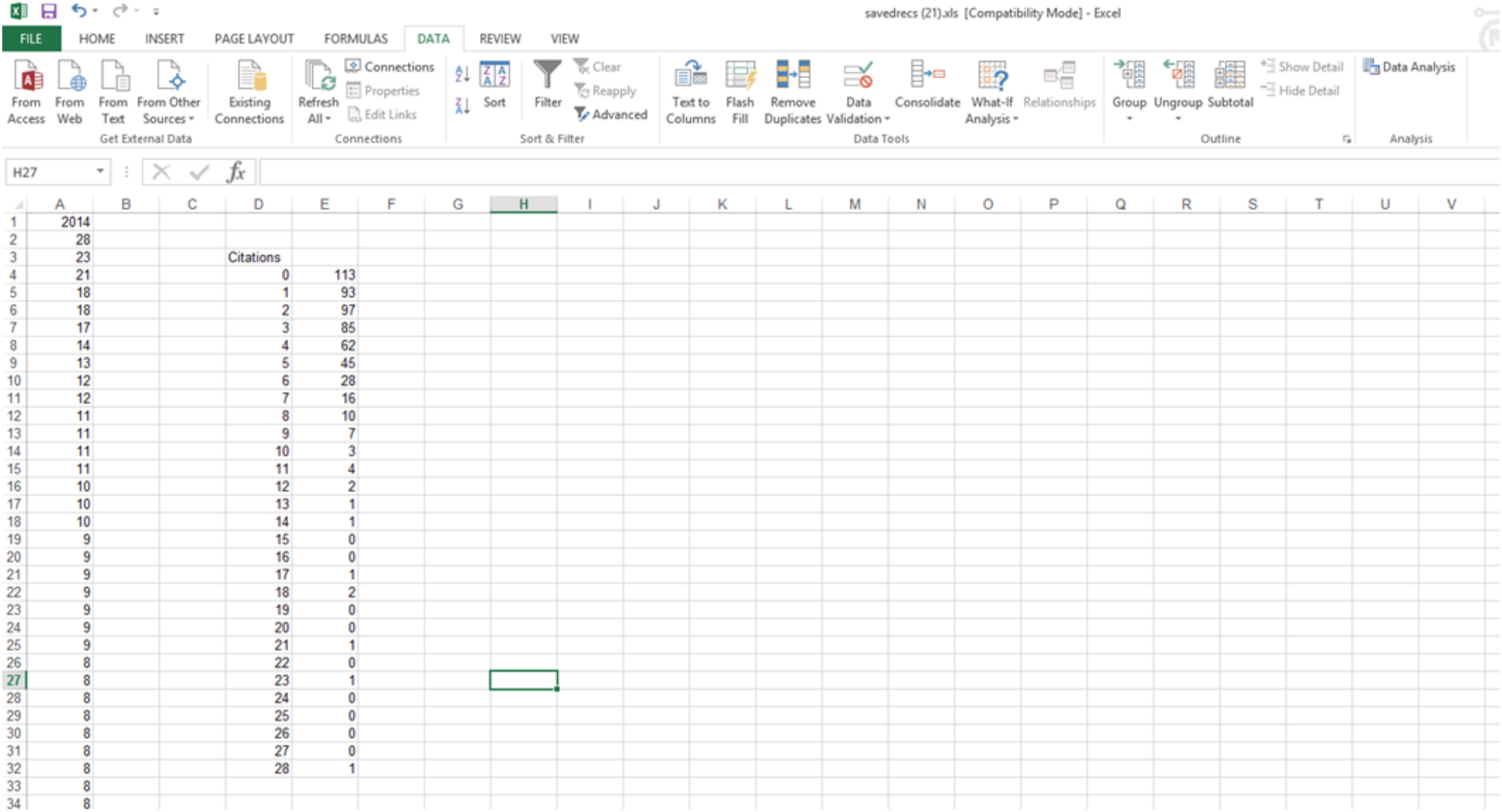

13. If you wish to determine the median, use Excel’s MEDIAN function on column A (excluding the 2014 label).

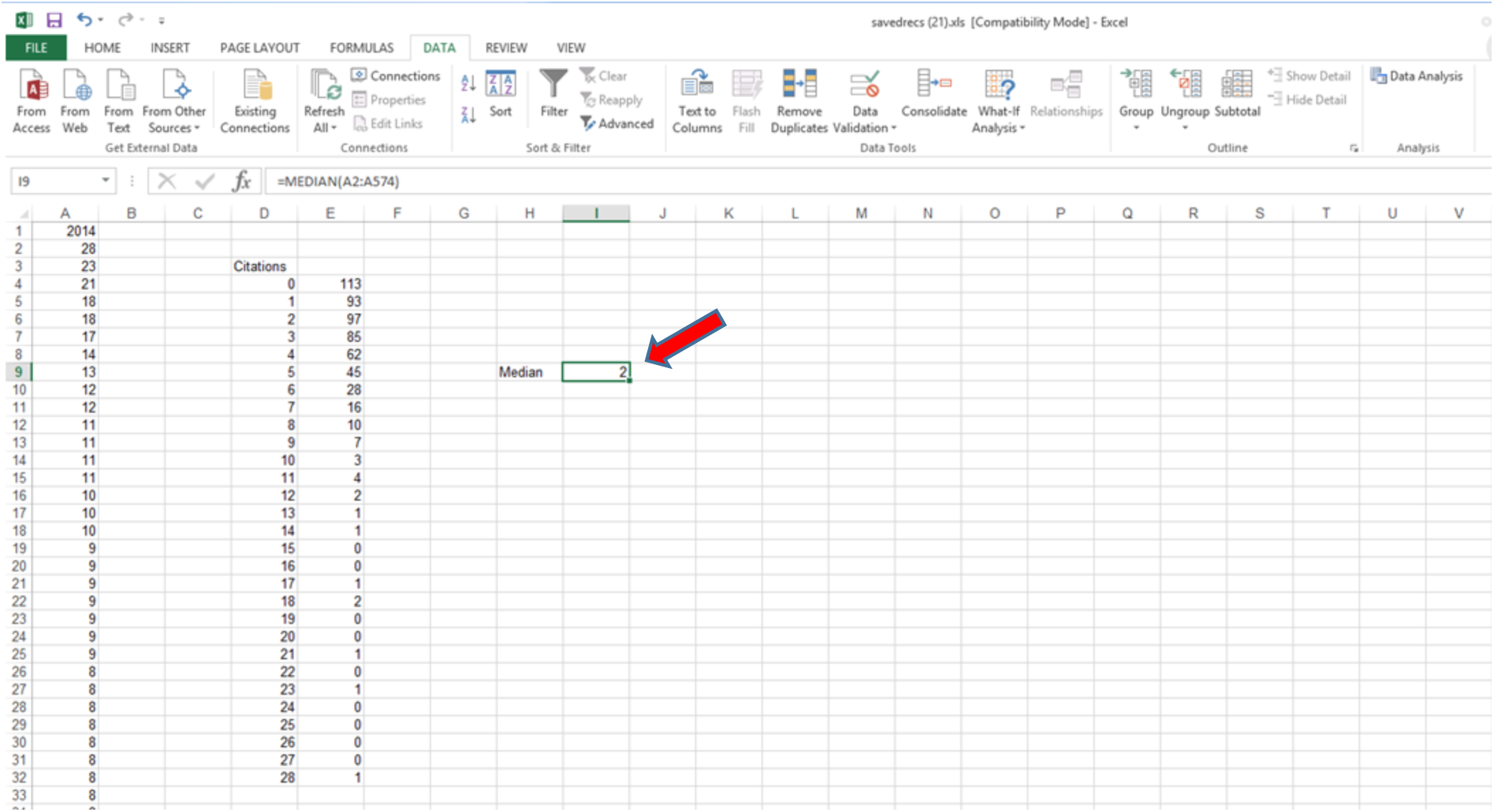

14. Then make a bar chart with the “Citations” field as the x-axis and the frequency counts as the y-axis. If desired, add vertical lines to indicate the JIF and the Median.

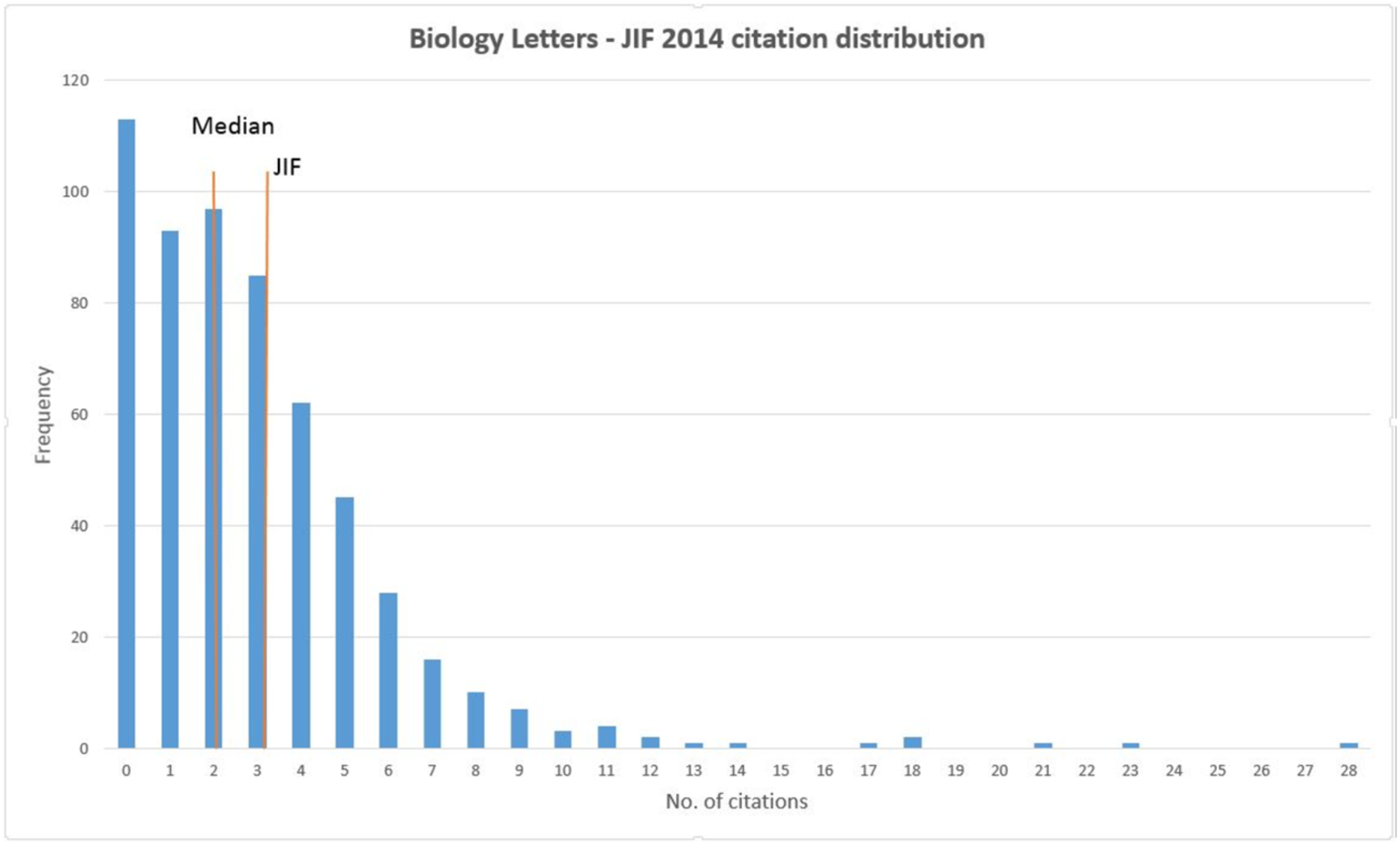

## Appendix 2 Method for generating the journal citation distribution graph from Scopus^™^ (2014 Impact Factor set)

The example given below is for generating distributions over the two-year window (2012-2013) that is used in calculation of the 2014 Journal Impact Factor. For later years, the two-year window should be adjusted accordingly.

1. In Scopus^™^, search for the journal using the ‘Source Title’ field and select the date range 2012-2013. Journal editors should check the resulting hit-list against the journal’s own record as tests showed that search results can return different article counts - largely due to duplicate records in the database (see 8. below). Users without access to internal records can check article counts via tables of contents.

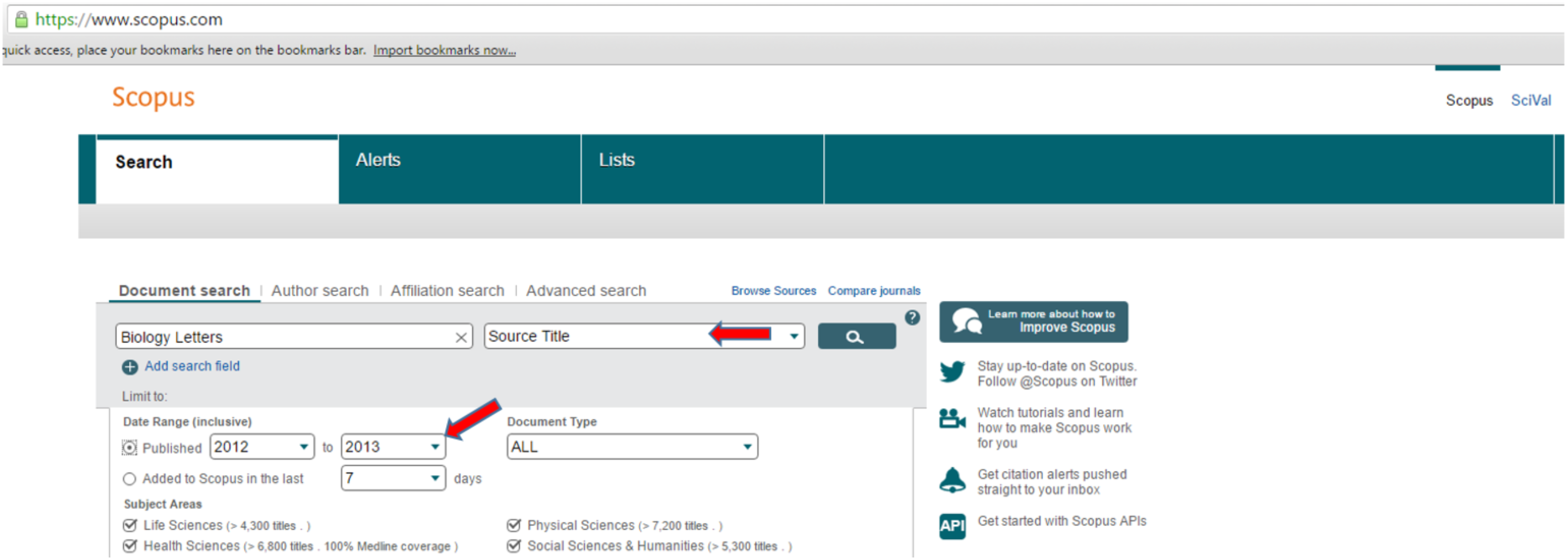

2. “Select all” from the resulting hit-list. (To match as closely as possible the distributions shown in the analyses in this paper, limit the document types in the search to ‘Articles’ and ‘Reviews’ using the buttons on the left hand side of the screen.)

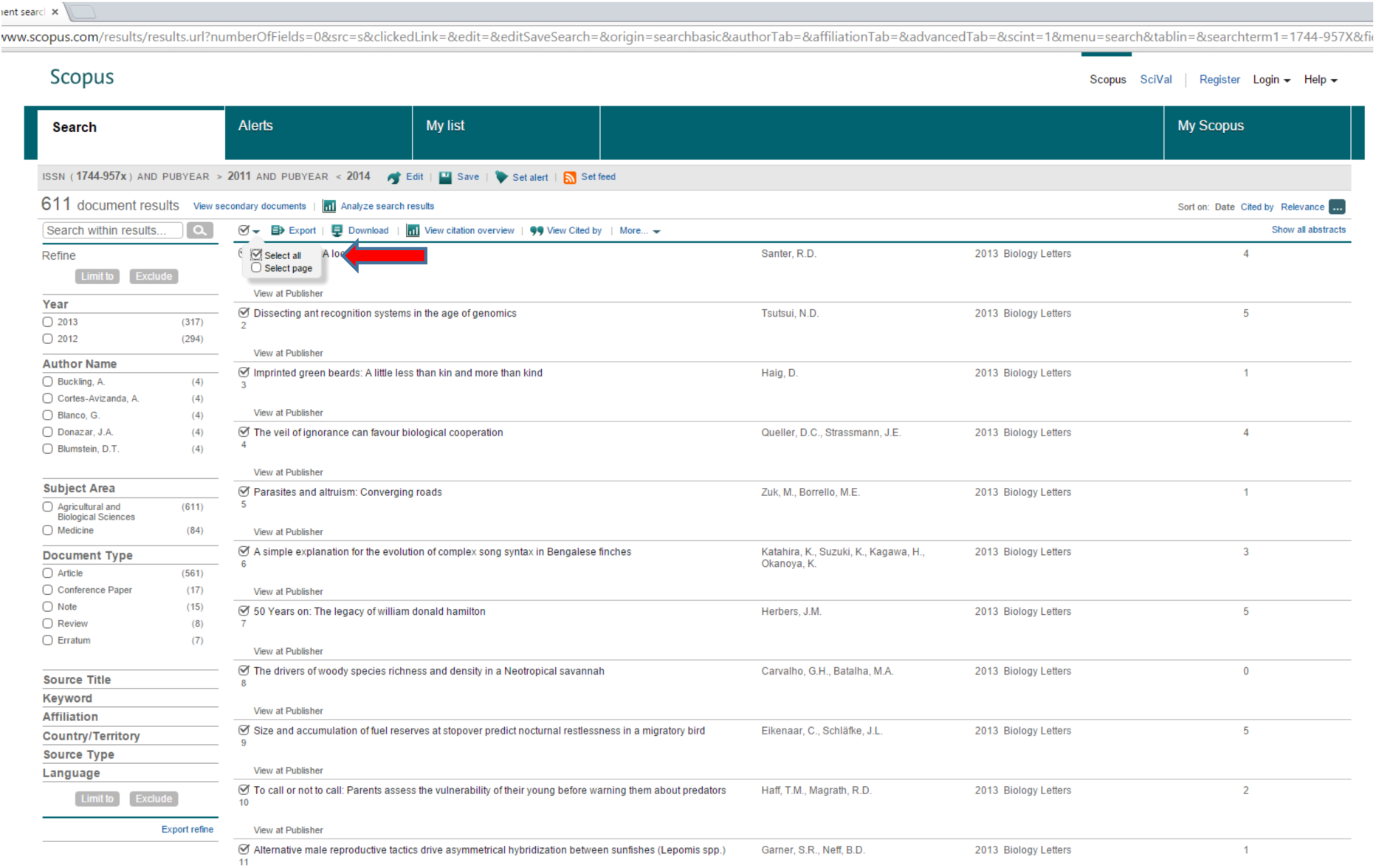

3. Click “view citation overview”.

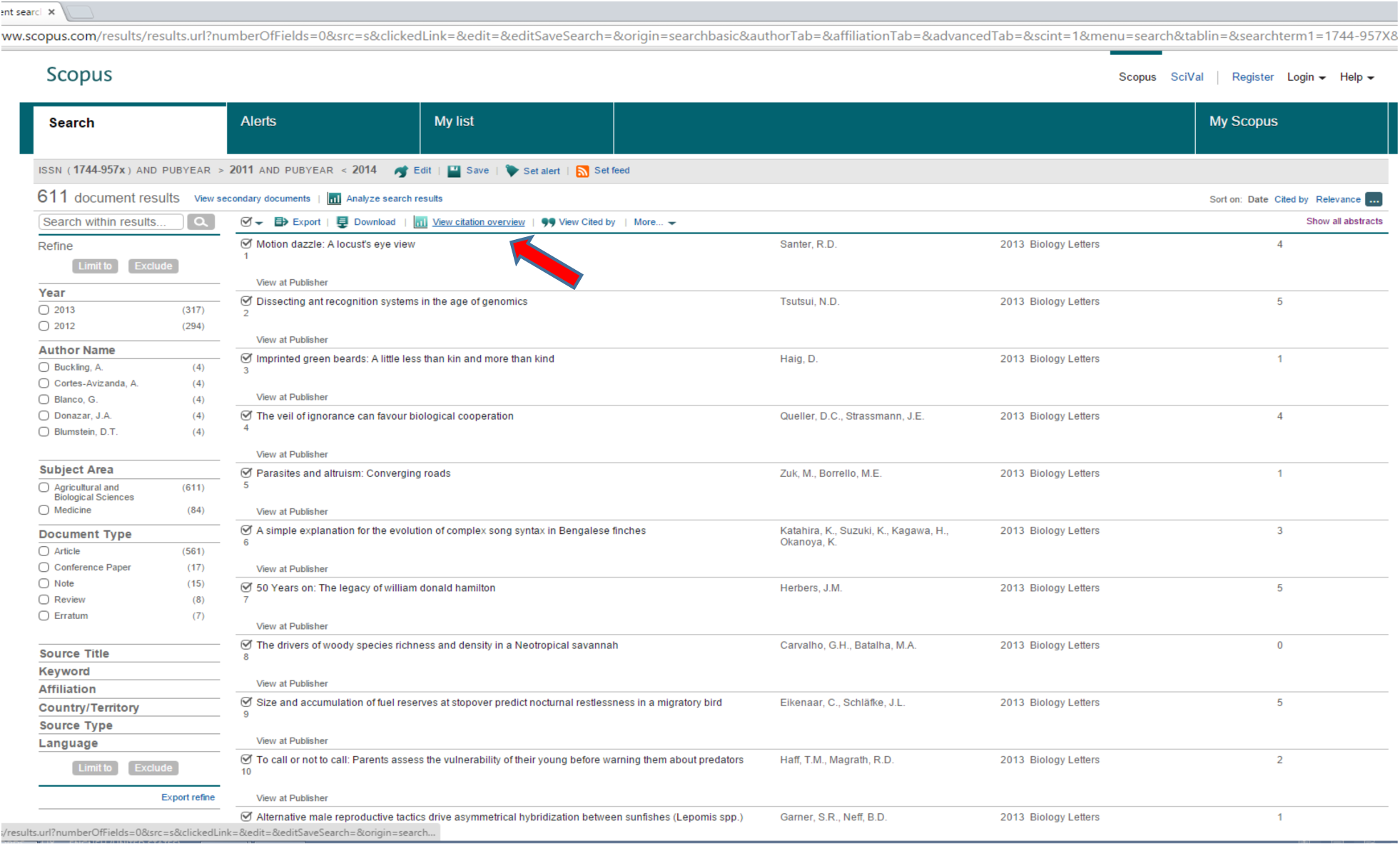

4. The Citation Overview will look something like this:

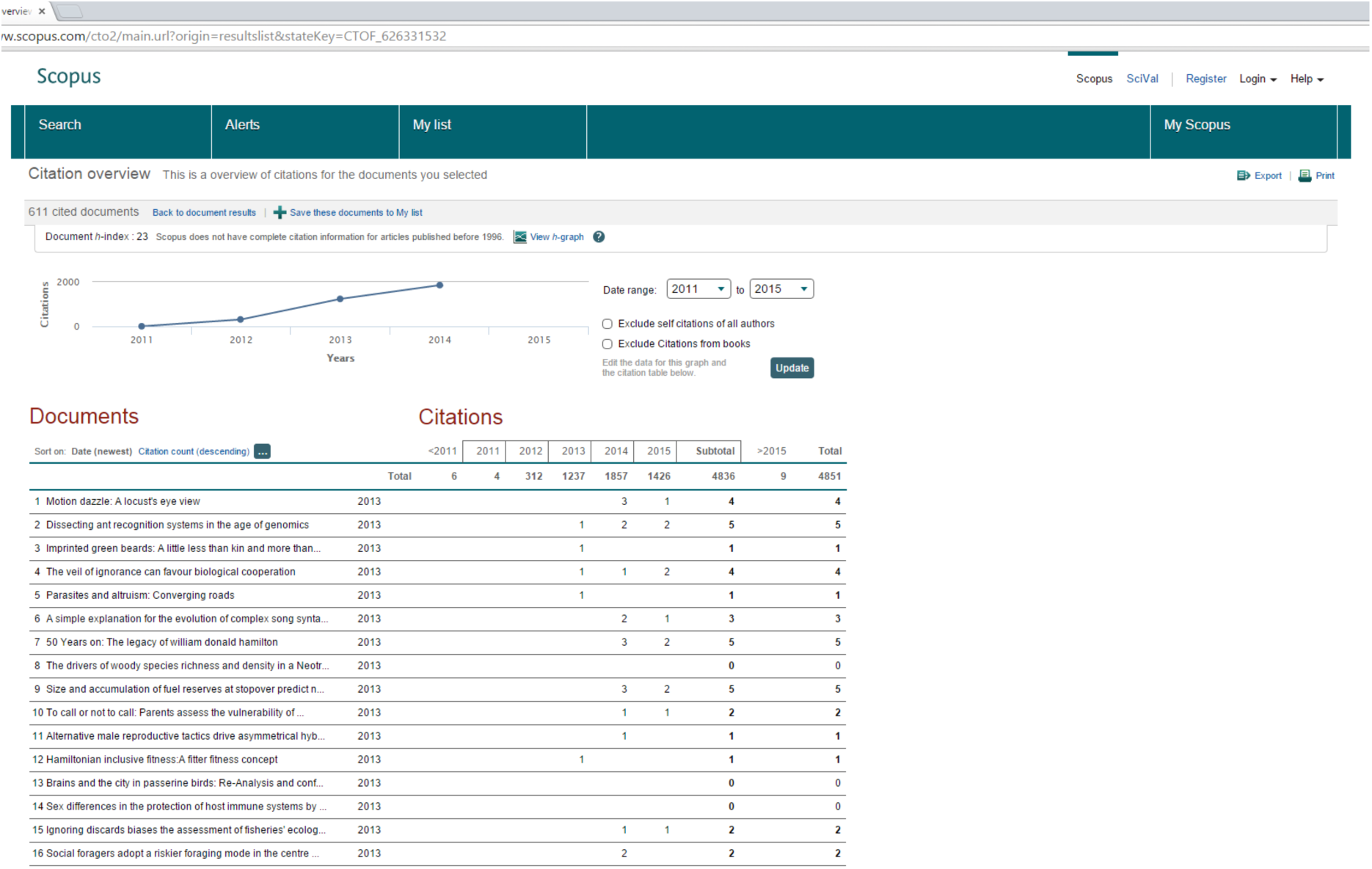

5. Select the date range 2014 (to get only citations in 2014) and click “update”.

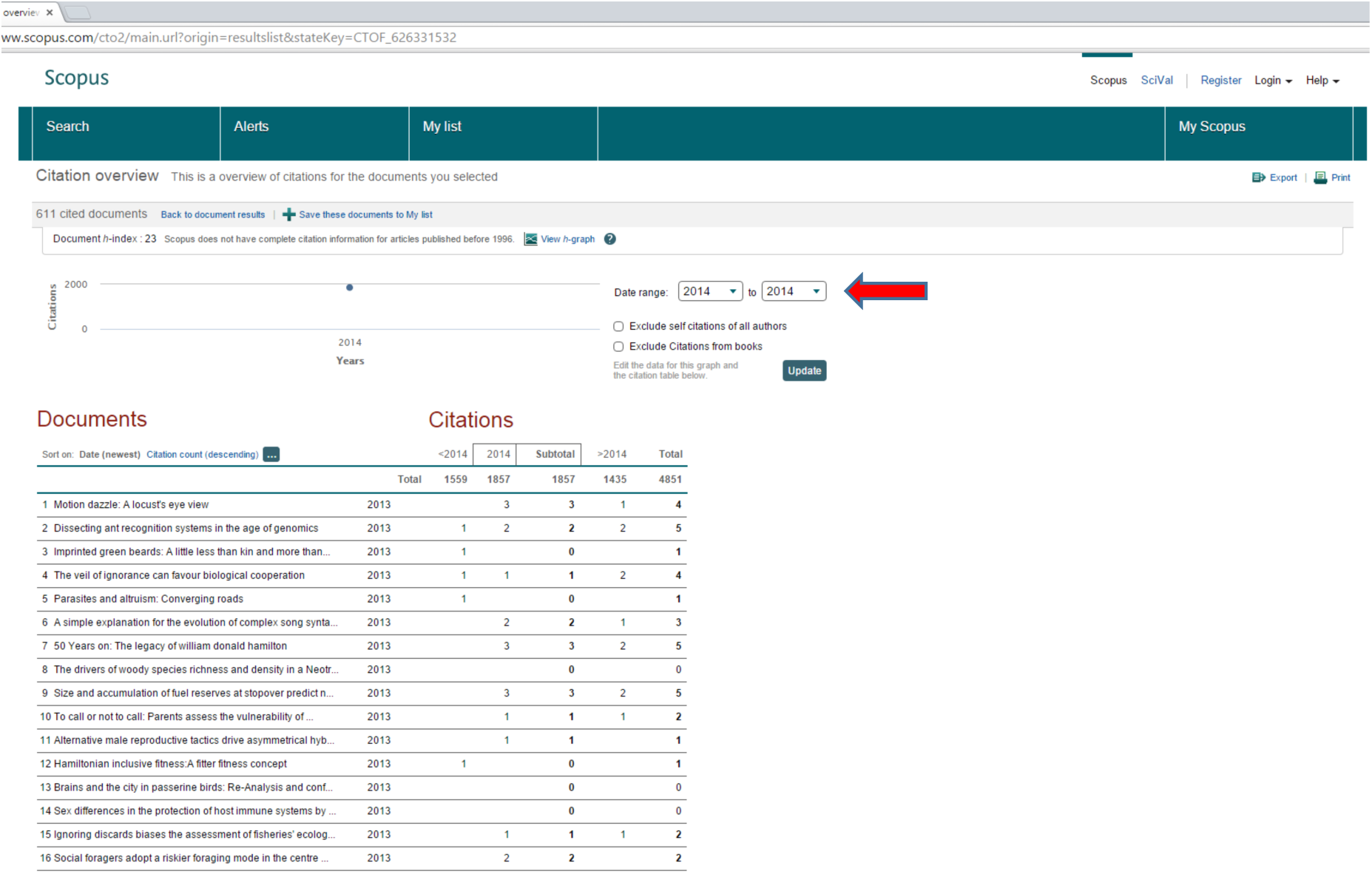

6. Then click “Export”.

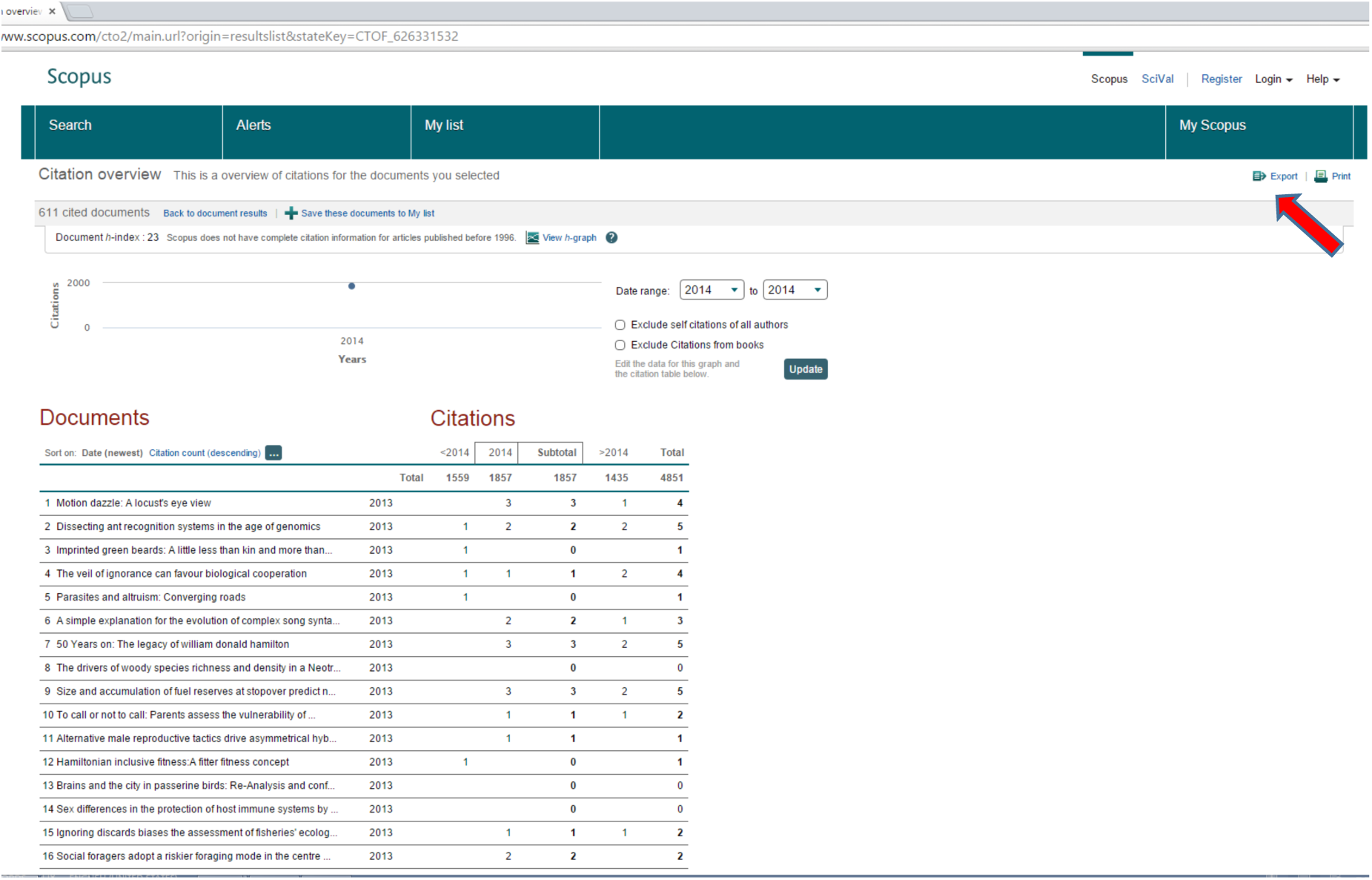

7. This will download a CSV (comma-separated values) file. Open it in Excel.

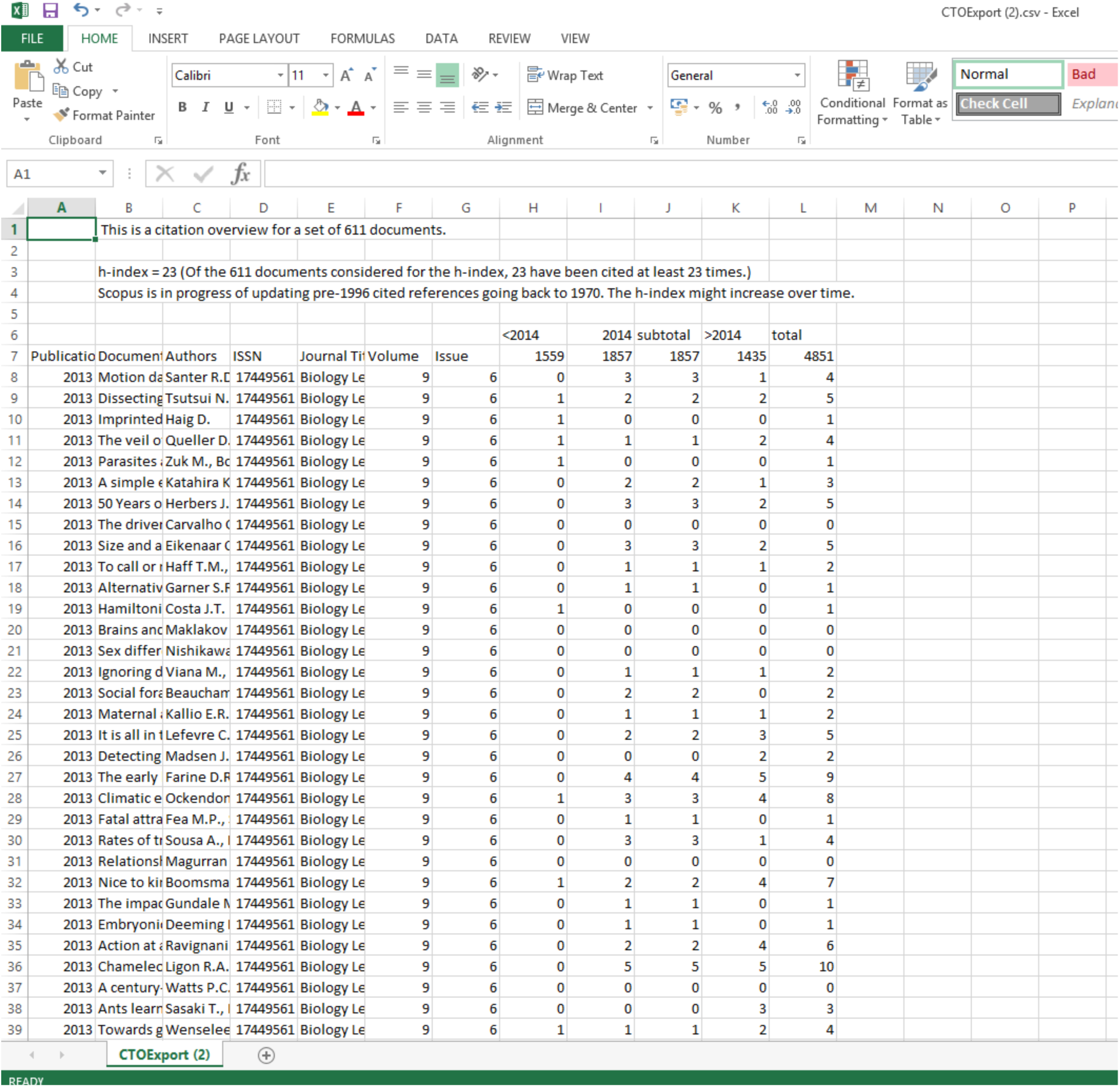

8. Scopus^™^ searches often produce duplicate records for the same paper both of which have associated citations. To resolve this, sort the records on the title column (A-Z) to make it easy to identify duplicates. For each pair, delete one, but make sure to add its citation count (*e.g*. in the 2014 column) to the remaining one to produce the correct total.

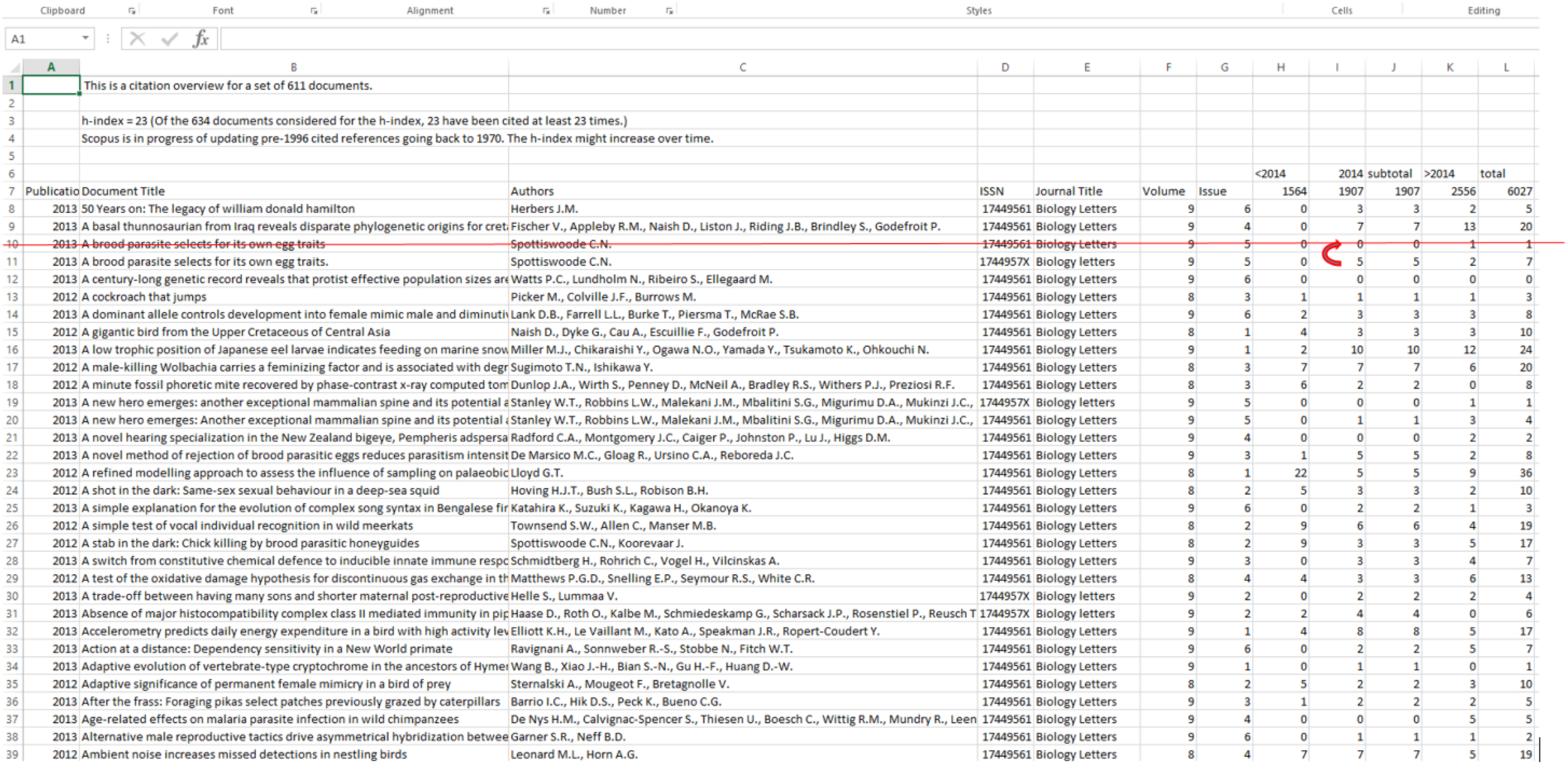

9. After de-duplication of the data, select the column for the citations received in 2014; (the other columns can be deleted).

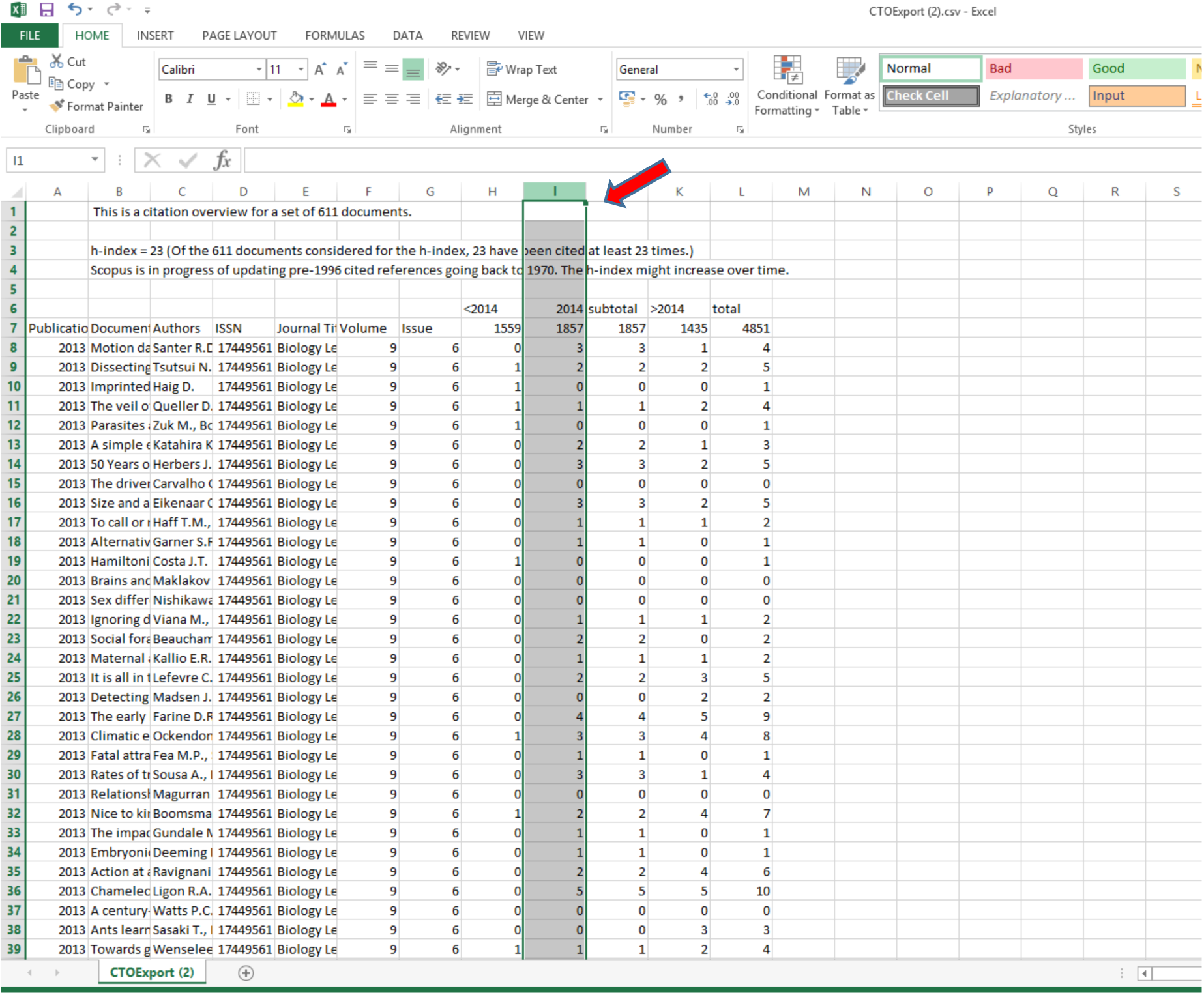

10. Sort the column into descending order – make sure to omit the row labels.

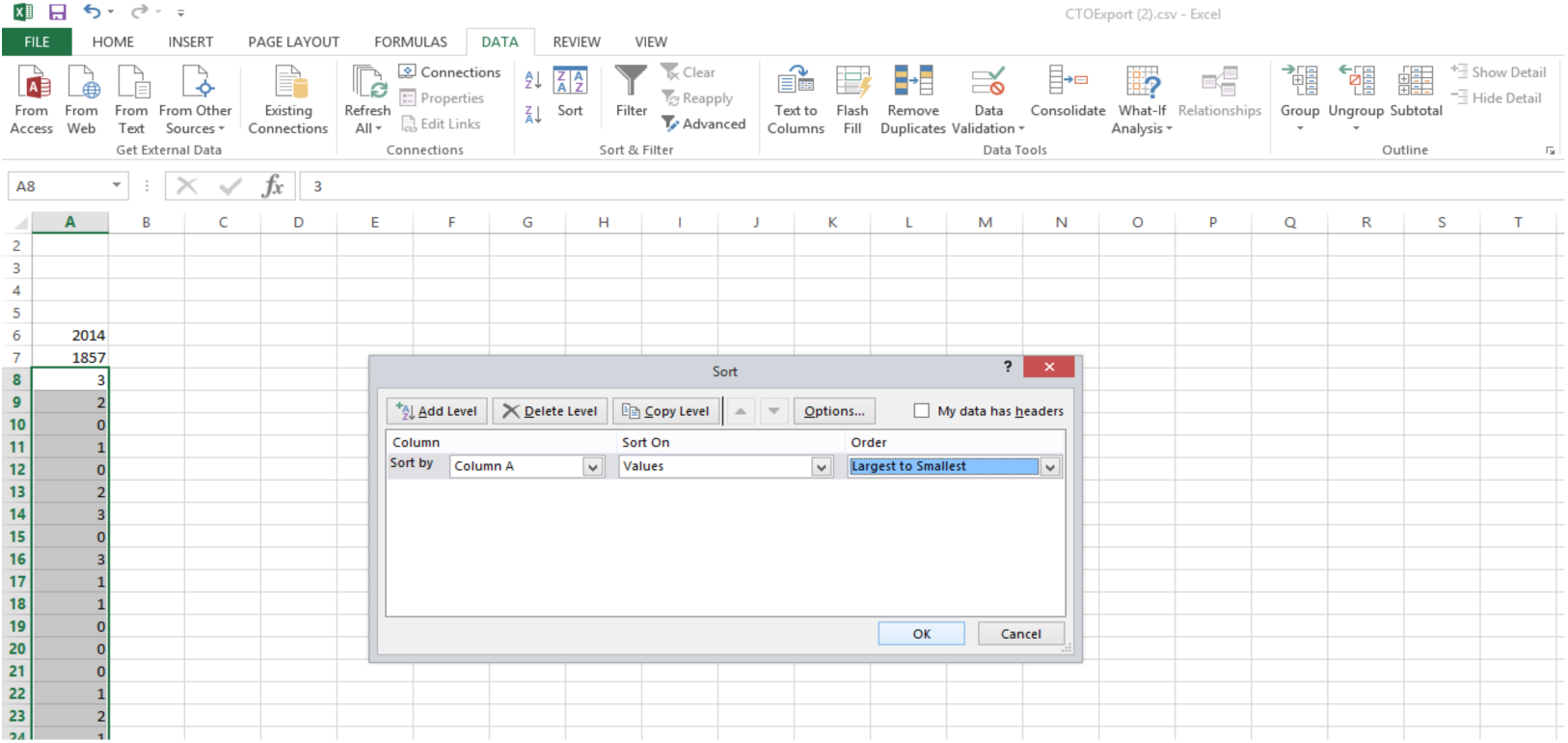

11. Note the maximum citation (x) count and create a new column containing 0 to X called “Citations”. In the example shown below, x = 28.

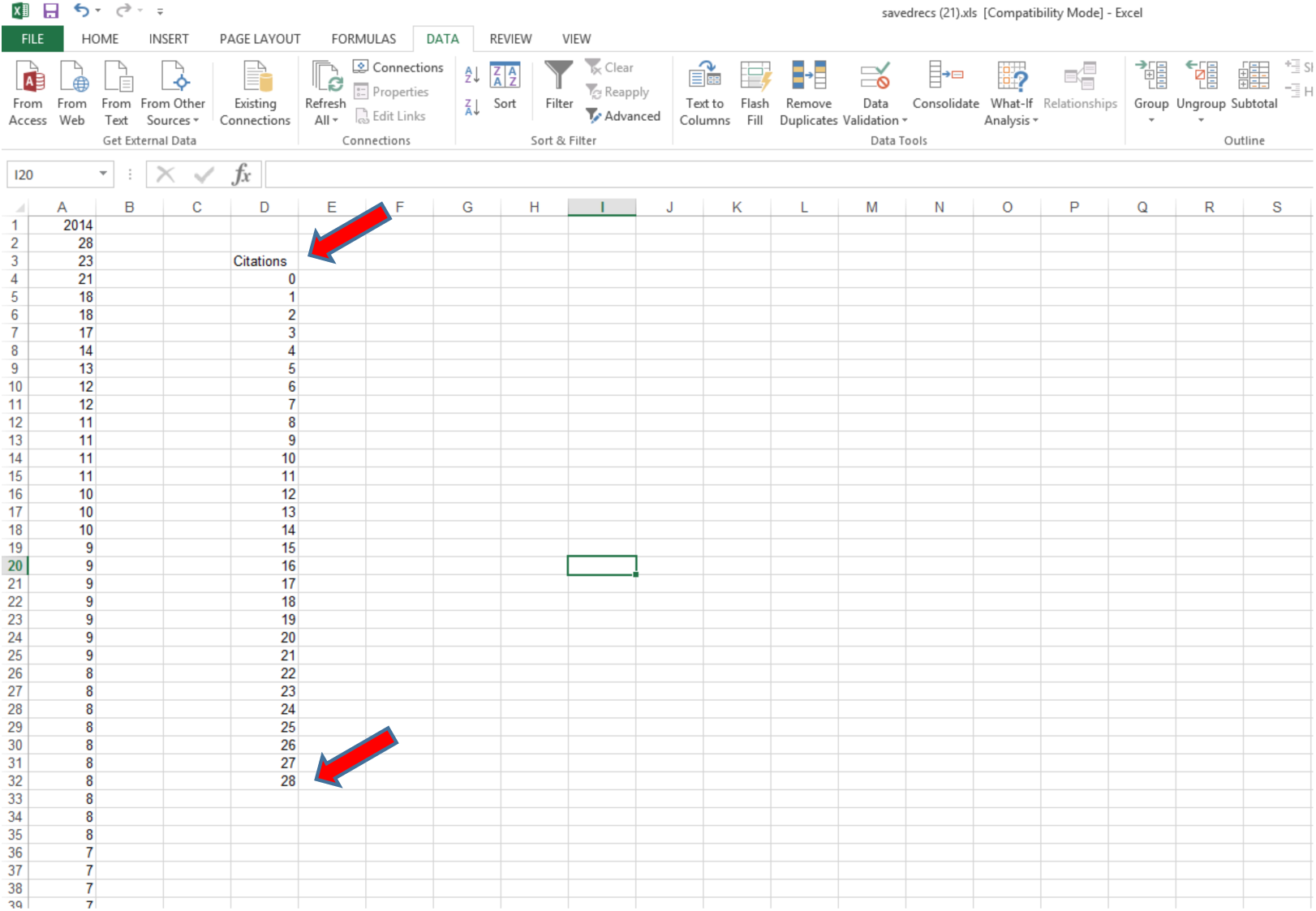

12. Enter the formula =COUNTIF(A:A,D4) into the cell next to the 0 citations (where A is the column containing the citations, and D4 is the cell with the zero citation count).

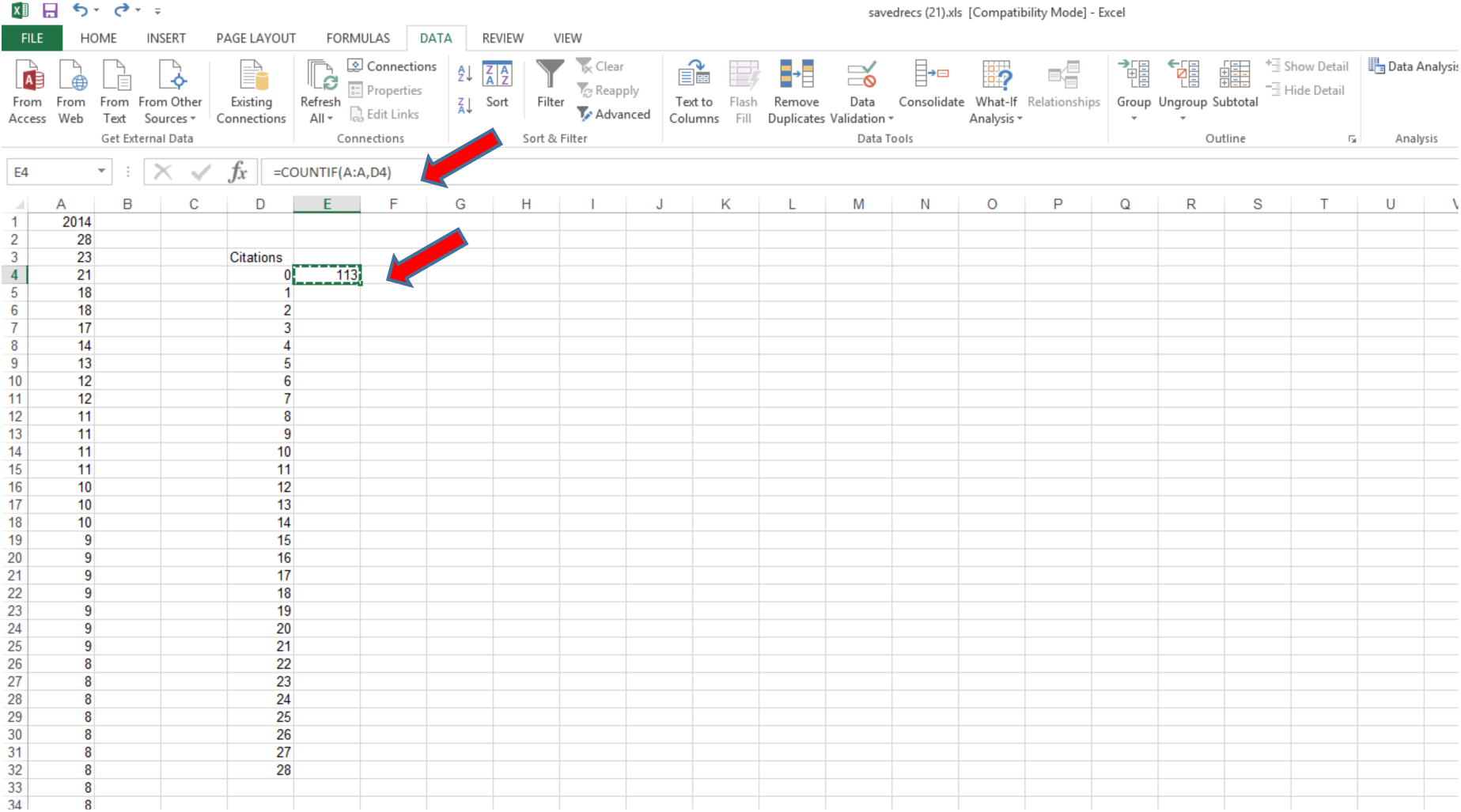

13. Copy and paste this formula into the remaining cells in the Citations column to generate the frequency distribution data.

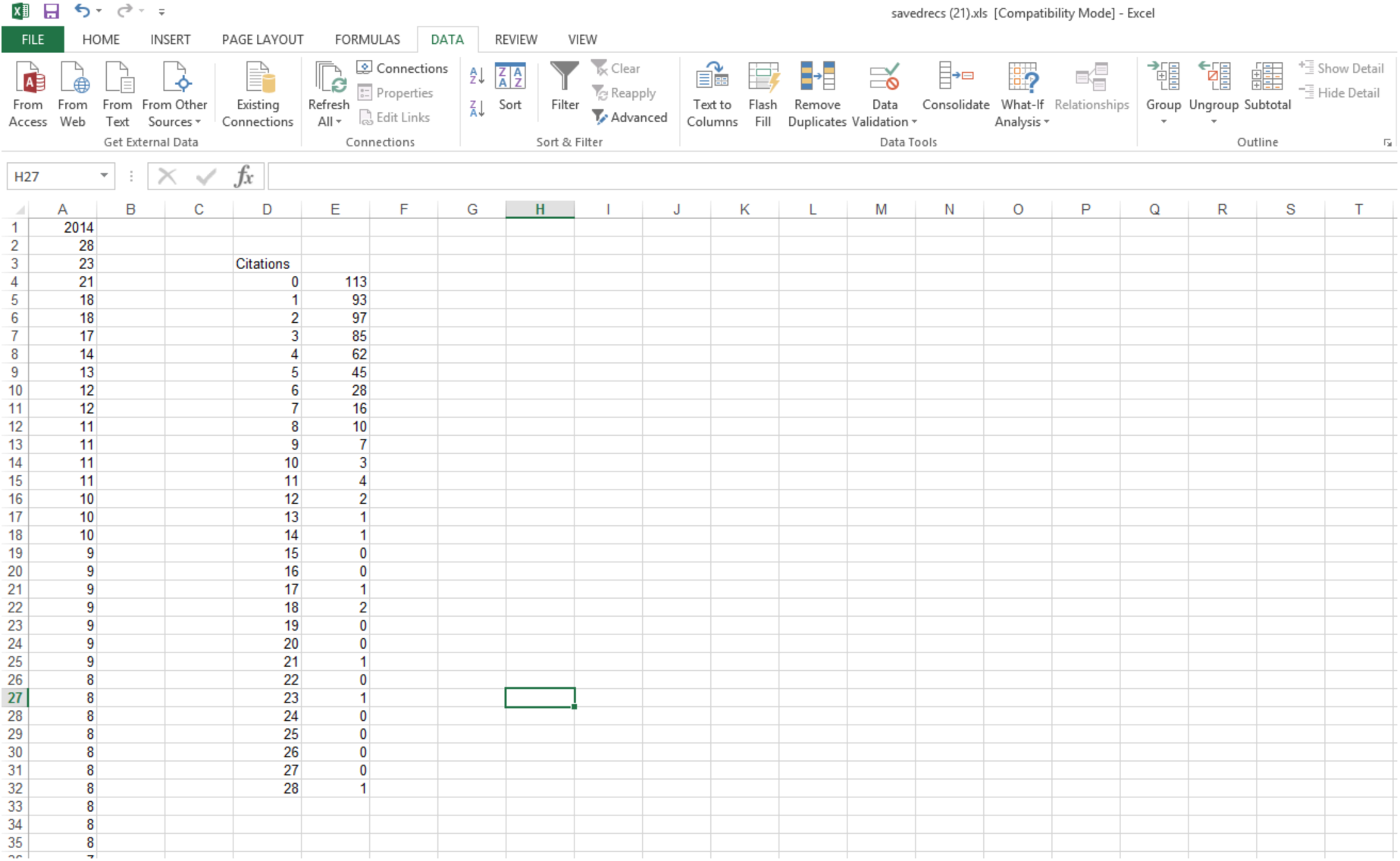

14. If you wish to determine the median, use Excel’s MEDIAN function on column A; be careful not to include the ‘2014’ label.

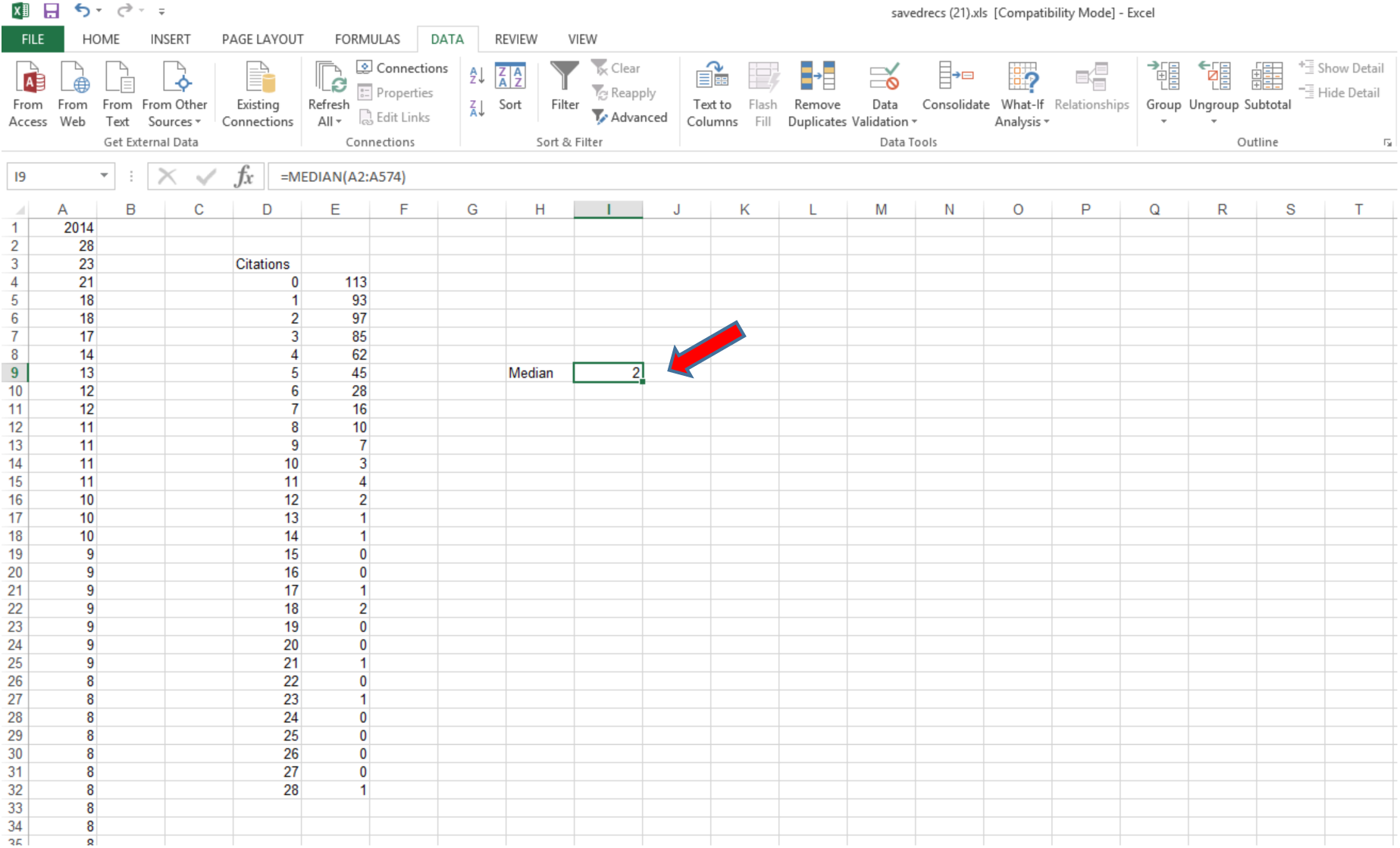

15. Then make a bar chart with the “Citations” field as the x-axis. If desired, add a vertical line to denote the JIF and the Median.

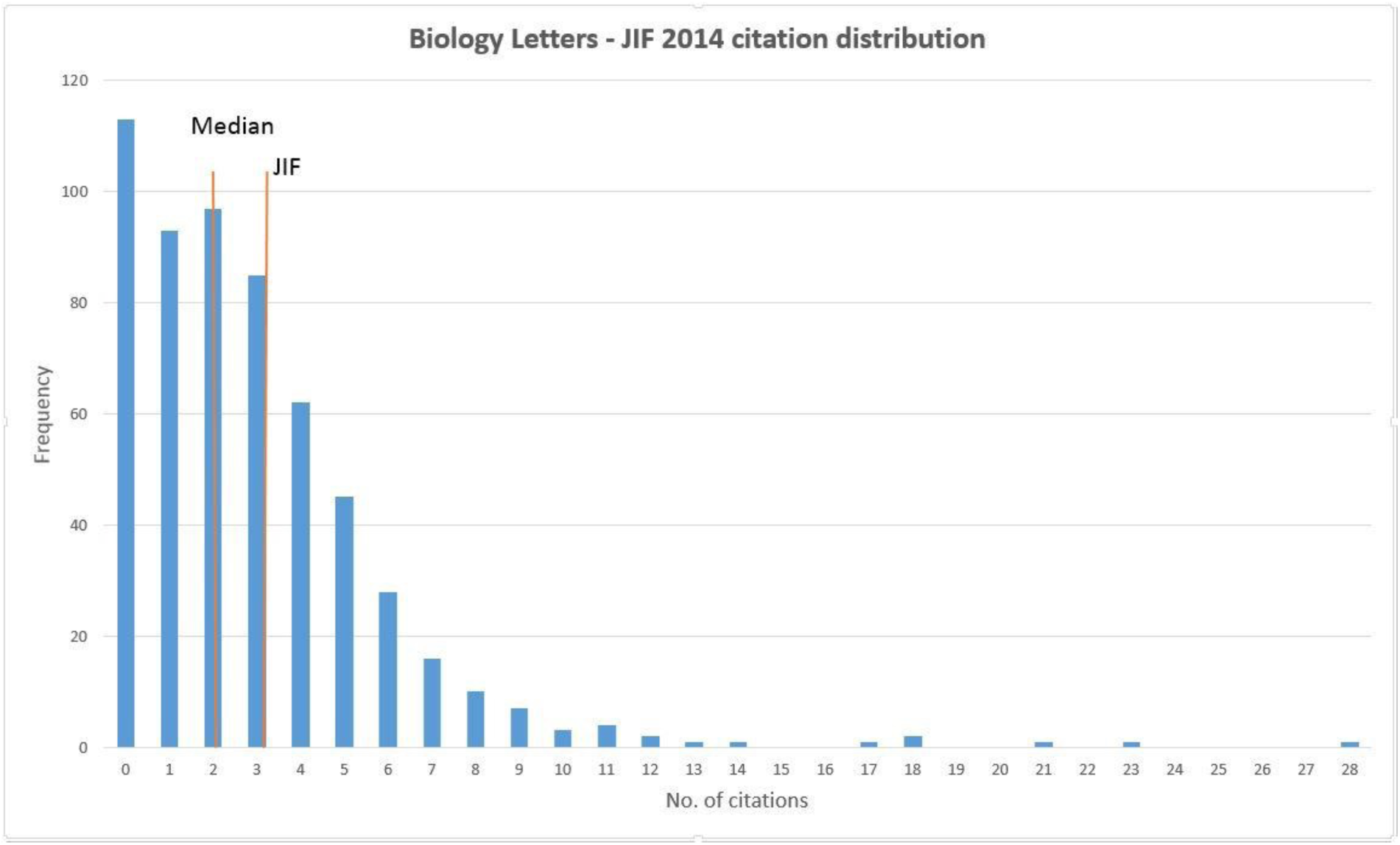

